# A cross-kingdom conserved ER-phagy receptor maintains endoplasmic reticulum homeostasis during stress

**DOI:** 10.1101/2020.03.18.995316

**Authors:** Madlen Stephani, Lorenzo Picchianti, Alexander Gajic, Rebecca Beveridge, Emilio Skarwan, Victor Sanchez de Medina Hernandez, Azadeh Mohseni, Marion Clavel, Yonglung Zeng, Christin Naumann, Mateusz Matuszkiewicz, Eleonora Turco, Christian Loefke, Baiying Li, Gerhard Durnberger, Michael Schutzbier, Hsiao Tieh Chen, Alibek Abdrakhmanov, Adriana Savova, Khong-Sam Chia, Armin Djamei, Irene Schaffner, Steffen Abel, Liwen Jiang, Karl Mechtler, Fumiyo Ikeda, Sascha Martens, Tim Clausen, Yasin Dagdas

## Abstract

Eukaryotes have evolved various quality control mechanisms to promote proteostasis in the ER. Selective removal of certain ER domains via autophagy (termed as ER-phagy) has emerged as a major quality control mechanism. However, the degree to which ER-phagy is employed by other branches of ER-quality control remains largely elusive. Here, we identify a cytosolic protein, C53, that is specifically recruited to autophagosomes during ER-stress, in both plant and mammalian cells. C53 interacts with ATG8 via a distinct binding epitope, featuring a shuffled ATG8 interacting motif (sAIM). C53 senses proteotoxic stress in the ER lumen by forming a tripartite receptor complex with the ER-associated ufmylation ligase UFL1 and its membrane adaptor DDRGK1. The C53/UFL1/DDRGK1 receptor complex is activated by stalled ribosomes and induces the degradation of internal or passenger proteins in the ER. Consistently, the C53 receptor complex and ufmylation mutants are highly susceptible to ER stress. Thus, C53 forms an ancient quality control pathway that bridges selective autophagy with ribosome-associated quality control at the ER.

## Introduction

Autophagy is an intracellular degradation process where eukaryotic cells remove harmful or unwanted cytoplasmic contents to maintain cellular homeostasis (Dikic and Elazar, 2018; Klionsky *et al*., 2011; Marshall and Vierstra, 2018). Recent studies have shown that autophagy is highly selective (Johansen and Lamark, 2019; Stolz *et al*., 2014), and is mediated by receptors that recruit specific cargo, such as damaged organelles or protein aggregates. Autophagy receptors and their cargo are incorporated into the growing phagophore through interaction with ATG8, a ubiquitin-like protein that is conjugated to the phagophore upon activation of autophagy (Stolz *et al*., 2014; Zaffagnini and Martens, 2016). The phagophore grows, and eventually forms a double-membrane vesicle termed the autophagosome. Autophagosomes then carry the autophagic cargo to lytic compartments for degradation and recycling. Selective autophagy receptors interact with ATG8 via conserved motifs called the ATG8 interacting motif (AIM) or LC3-interacting region (LIR) (Birgisdottir *et al*., 2013). In contrast to mammals and yeast, cargo receptors that mediate organelle recycling remains mostly elusive in plants (Stephani and Dagdas, 2019).

The endoplasmic reticulum (ER) is a highly dynamic heterogeneous cellular network that mediates folding and maturation of −40 % of the proteome (Sun and Brodsky, 2019; Walter and Ron, 2011). Proteins that pass through the ER include all secreted and plasma membrane proteins and majority of the organellar proteins. This implies, ER could handle up to a million client proteins in a cell every minute (Karagoz *et al*., 2019). Unfortunately, the folding process is inherently error prone and misfolded proteins are toxic to the cell (Fregno and Molinari, 2019; Karagoz *et al*., 2019; Sun and Brodsky, 2019). To maintain the proteostasis in the ER, eukaryotes have invested in quality control mechanisms that closely monitor, and if necessary, trigger the removal of terminally misfolded proteins. Degradation of the faulty proteins is mediated by proteasomal and vacuolar degradation pathways (Fregno and Molinari, 2019).

One of the main vacuolar/lysosomal degradation processes is ER-phagy. It has emerged as a major quality control pathway, and defects in ER-phagy is linked to various diseases (Chino and Mizushima, 2020; Hubner and Dikic, 2019; Stolz and Grumati, 2019; Wilkinson, 2019). ER-phagy involves cargo receptors that mediate removal of certain regions of the ER via autophagy. Several ER-resident ER-phagy receptors have been identified. These include Fam134B, RTN3L, ATL3, Sec62, CCPG1, and TEX264 in mammals and ATG39 and ATG40 in yeast (An *et al*., 2019; Chen *et al*., 2019; Chino *et al*., 2019; Fumagalli *et al*., 2016; Grumati *et al*., 2017; Khaminets *et al*., 2015; Mochida *et al*.; Smith *et al*., 2018). A recent study showed reticulon proteins could also function as ER-phagy receptors in plants (Zhang *et al*., 2020). These receptors are activated during starvation or stress conditions and remodel the ER network to maintain proteostasis. Despite the emerging links, how ER-phagy cross-talks with the rest of the ER quality control pathways needs further investigation (Chino and Mizushima, 2020; Dikic, 2018).

Here, using a peptide-competition coupled affinity proteomics screen, we identified a highly conserved cytosolic protein, C53, that is specifically recruited into autophagosomes during ER stress. C53 interacts with plant and mammalian ATG8 isoforms via non-canonical ATG8 interacting motifs (AIM), termed shuffled AIM (sAIM). C53 is recruited to the ER by forming a ternary receptor complex with the UFL1, the E3 ligase that mediates ufmylation, and its ER membrane adaptor DDRGK1 (Gerakis *et al*., 2019). C53 mediated autophagy is activated upon ribosome stalling during co-translational protein translocation and mediates degradation of model ribosome stalling constructs and various other ER resident or folded proteins. Consistently, C53 is crucial for ER stress tolerance in both plants and HeLa cells.

## Results

### C53 interacts with plant and mammalian ATG8 isoforms in an ER-stress dependent manner

To identify specific cargo receptors that mediate selective removal of ER compartments during proteotoxic stress, we performed an immunoprecipitation coupled to mass spectrometry (IP-MS) screen to identify AIM-dependent ATG8 interactions triggered by ER stress. We hypothesized that a synthetic AIM peptide that has higher affinity for ATG8 can outcompete, and thus reveal, AIM-dependent ATG8 interactors. To identify this synthetic peptide, we performed a peptide array analysis that revealed the AIM *wt* peptide (Figure S1a, b; Table S1). Using isothermal titration calorimetry (ITC), we showed that the AIM *wt* binds ATG8 with nanomolar affinity (*K*_D_=-700 nM), in contrast to the AIM mutant peptide (AIM *mut*), which does not show any binding (Figure S1c-f) or the low micromolar-range affinities measured for most cargo receptors (Zaffagnini and Martens, 2016). As plants have an expanded set of ATG8 proteins, we first tested if any of the ATG8 isoforms specifically responded to ER stress induced by tunicamycin (Kellner *et al*., 2016). Tunicamycin inhibits glycosylation and leads to proteotoxic stress at the ER (Bernales *et al*., 2006). Quantification of ATG8 puncta in transgenic seedlings expressing GFP-ATG8A-I revealed that tunicamycin treatment significantly induced all nine ATG8 isoforms (Figure S2). Since all ATG8 isoforms were induced, we chose ATG8A, and performed peptide competition coupled IP-MS analysis (See methods for detailed description). In addition to well-known AIM dependent ATG8 interactors such as ATG4 (Autophagy related gene 4) and NBR1 (Neighbour of BRCA1) (Wild *et al*., 2014), our analyses revealed that the highly conserved cytosolic protein C53 (aliases: CDK5RAP3, LZAP, IC53, HSF-27) is an AIM-dependent ATG8 interactor (Figure 1a, Table S2, Figure S3).

**Figure 1.**
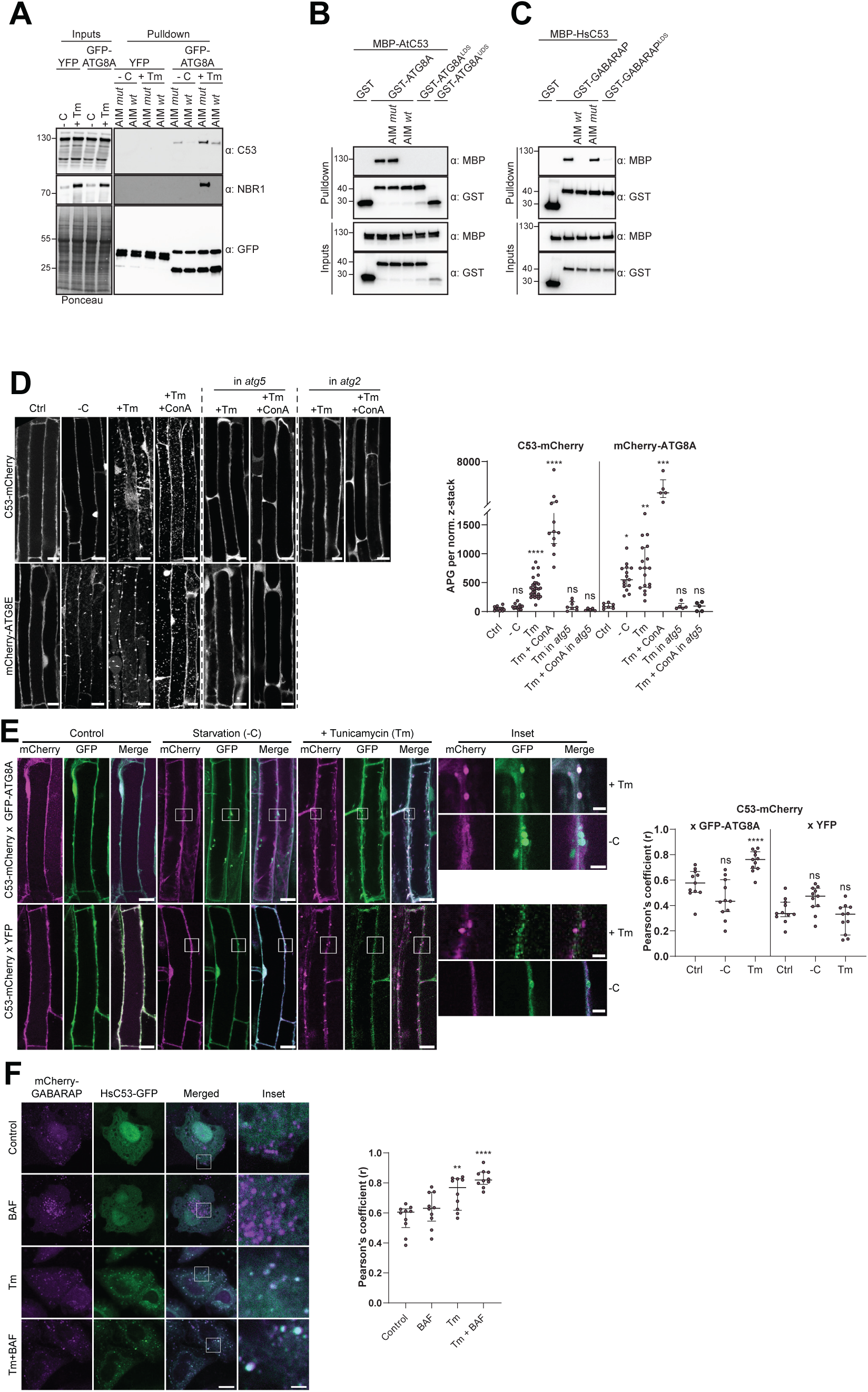
C53 binds ATG8 in an AIM dependent manner and undergoes ER-stress specific autophagic degradation. (**A**) Peptide competition coupled *in vivo* pull-down revealed C53 as an AIM dependent ATG8 interactor. */n vivo* co-immunoprecipitation of extracts of Arabidopsis seedlings expressing YFP alone or GFP-ATG8A incubated in sucrose (-C)-deficient or 10 µg/ml tunicamycin (Tm) containing media. The peptides AIM *wt* and AIM *mut* were added to a final concentration of 200 µM. Input and bound proteins were visualized by immunoblotting with anti-GFP, anti-NBR1, and anti-C53 antibodies. (**B, C**) **AtC53 and HsC53 interact with ATG8A and GABARAP, respectively, in an AIM-dependent manner**. Bacterial lysates containing recombinant protein were mixed and pulled down with glutathione magnetic agarose beads. The peptides AIM *wt* and AIM *mut* were added to a final concentration of 200 µM. Input and bound proteins were visualized by immunoblotting with anti-GST and anti-MBP antibodies. LDS=LIR-Docking-Site mutant (Marshall *et al*., 2019); UDS = Ubiquitin Docking Site mutant (Marshall *et al*., 2019). (**D**) AtC53 is specifically recruited to puncta upon ER stress and this depends on ATG5 and ATG2. *Left Panel*, representative confocal images of transgenic Arabidopsis seedlings expressing C53-mCherry and mCherry-ATG8E in Col-0 wild type, *atg5* and *atg2* mutant backgrounds. 6-day old seedlings were incubated in either control, sucrose (-C)-deficient, tunicamycin (10 µg/ml), or tunicamycin (Tm, 10 µg/ml) + Concanamycin (ConA, 1 µM) containing media. Scale bars, 10 µm. *Right Panel*, Quantification of the autophagosomes (APG) per normalized Z-stacks. Bars represent the mean (± SD) of at least 10 biological replicates. (**E**) **AtC53 puncta colocalize with GFP-ATG8A labelled autophagosomes during ER stress**. *Left Panel*, Co-localization analyses of single plane confocal images obtained from transgenic Arabidopsis roots co-expressing C53-mCherry (magenta) with GFP-ATG8A or YFP alone (green). 4-day old seedlings were incubated in either control, sucrose deficient (-C), or tunicamycin containing media. Scale bars, 20 µm. Inset scale bars, 2 µm. *Right Panel*, Pearson’s Coefficient (r) analysis of the colocalization of C53-mCherry with GFP-ATG8A or YFP alone. Bars represent the mean (± SD) of at least 5 biological replicates. (**F**) HsC53 puncta colocalize with mCherry-GABARAP labelled autophagosomes during ER stress. *Left Panel*, Confocal images of PFA fixed HeLa cells transiently expressing C53-GFP (green) and mCherry-GABARAP (magenta). Cells were either untreated (Control) or treated with tunicamycin (Tm) or Tm + Bafilomycin (BAF). Scale bar, 20 µm. Inset scale bar, 2 µm. *Right Panel*, Pearson’s Coefficient analysis of the colocalization of HsC53-GFP with mCherry-GABARAP under control and Tm treated conditions. Bars represent the mean (± SD) of at least 5 biological replicates. Significant differences are indicated with * when p value ≤ 0.05, ** when p value ≤ 0.01, and *** when p value ≤ 0.001.

To confirm our IP-MS results, we performed *in vitro* pull-down experiments. *Arabidopsis thaliana* (At) C53 specifically interacted with GST-ATG8A, and this interaction was outcompeted with the AIM *wt*, but not AIM *mut* peptide. Consistently, ATG8 receptor accommodating site mutations (LDS − LIR Docking Site) prevented C53 binding (Figure 1b). We extended our analysis to all Arabidopsis ATG8 isoforms and showed that AtC53 interacts with 8 of 9 Arabidopsis isoforms. To probe for evolutionary conservation of C53-ATG8 interaction, we tested the orthologous proteins from the basal land plant *Marchantia polymorpha* (Mp) and showed that MpC53 interacts with 1 of 2 Marchantia ATG8 isoforms (Figure S4a, b). As C53 is highly conserved in multicellular eukaryotes and has not been characterized as an ATG8 interactor in mammals, we tested whether human C53 (HsC53) interacts with human ATG8 isoforms (LC3A-C, GABARAP, -L1, -L2). HsC53 interacted with GABARAP and GABARAPL1 in an AIM-dependent manner via the LIR docking site, similar to plant C53 homologs (Figure 1c, Figure S4c, d). Together, these data suggest that C53-ATG8 interaction is conserved across kingdoms.

In order to examine the *in vivo* link between C53 and ATG8 function, we generated transgenic AtC53-mCherry Arabidopsis lines and measured autophagic flux during ER stress. Without stress, AtC53 displayed a diffuse cytosolic pattern. Similarly, upon carbon starvation (-C, Figure 1d), which is commonly used to activate bulk autophagy, AtC53-mCherry remained mostly diffuse (Marshall and Vierstra, 2018). However, tunicamycin (Tm) treatment led to a rapid increase in AtC53 puncta. The number of puncta was further increased upon concanamycin A (ConA) treatment that inhibits vacuolar degradation, suggesting that AtC53 puncta are destined for vacuoles (Figure 1d). The AtC53 puncta disappeared when AtC53-mCherry lines were crossed into core autophagy mutants *atg5* and *atg2*, confirming that formation of AtC53 puncta is dependent on macroautophagy (Figure 1d). Consistent with this, other ER-stressors such as phosphate starvation, cyclopiazonic acid (CPA), and dithiothreitol (DTT) treatments also induced AtC53 puncta (Figure S5) (Fumagalli *et al*., 2016; Naumann *et al*., 2018; Smith *et al*., 2018). The AtC53 puncta co-localized with GFP-ATG8A and GFP-ATG11, indicating that they are autophagosomes (Figure 1e, Figure S6a). Moreover, as recently shown for other selective autophagy receptors, AtC53 and HsC53 directly interacted with the mammalian ATG11 homolog FIP200 (PTK2/FAK family-interacting protein of 200 kDa) (Figure S6b) (Lahiri and Klionsky, 2018; Ravenhill *et al*., 2019; Turco *et al*., 2019; Vargas *et al*., 2019). Ultrastructural analysis using immunogold labelling also confirmed localization of AtC53 at autophagosomes during ER-stress (Figure S7). Similar to plant proteins, transfected HsC53-GFP co-localized with mCherry-GABARAP upon tunicamycin treatment in HeLa cells. The number of HsC53 puncta increased upon bafilomycin (BAF) treatment, which inhibits lysosomal degradation; suggesting that HsC53 puncta eventually fuse with lysosomes (Figure 1f). To support our imaging-based autophagic flux assays, we also performed western blot based autophagic flux analyses, which further demonstrated ER-stress specific autophagic degradation of AtC53 and HsC53 (Figure 2).

**Figure 2.**
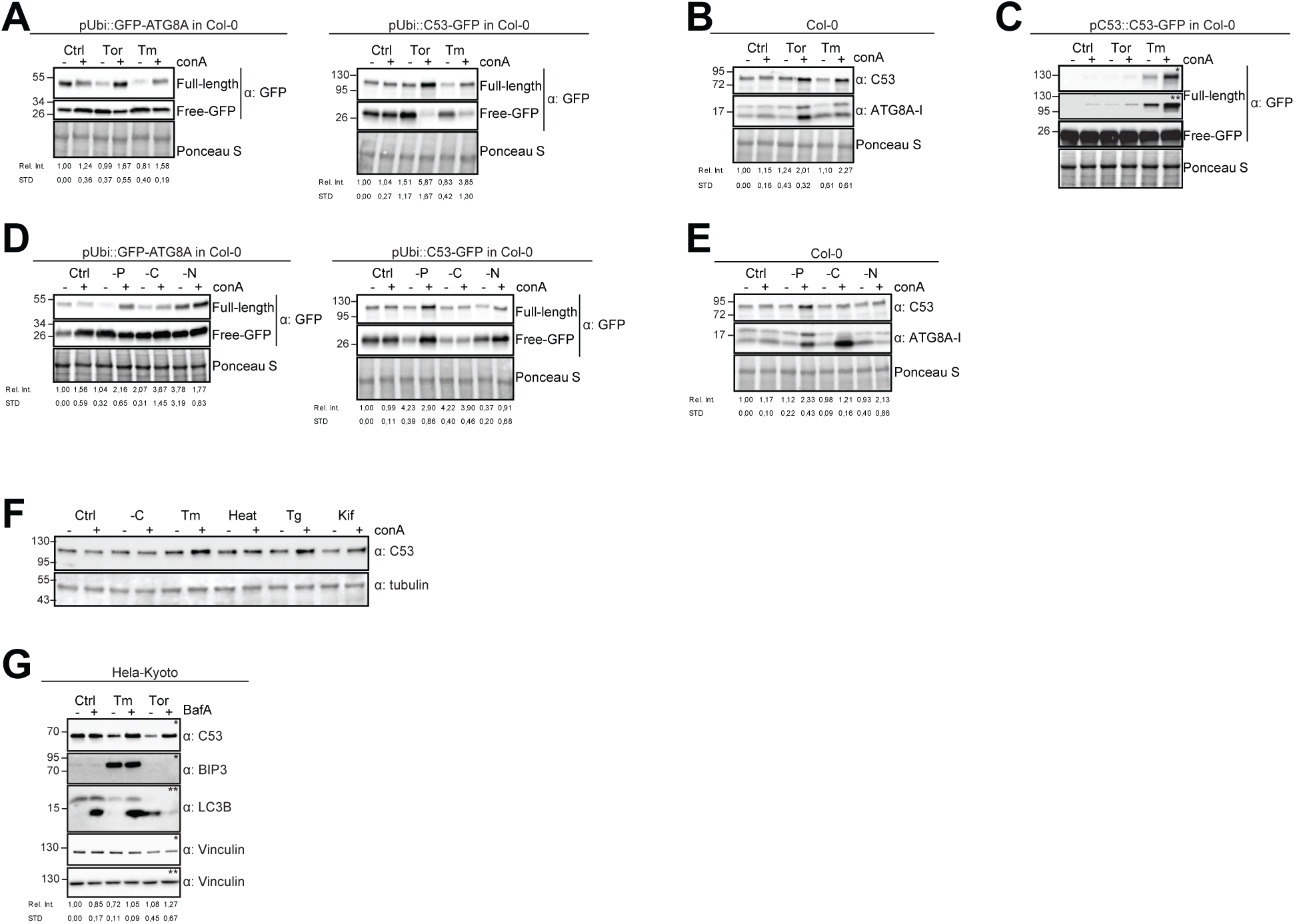
Autophagic flux analysis of AtC53 and HsC53 show that C53 autophagic flux is induced during ER stress. (A-C) AtC53 flux is induced by Torin and tunicamycin treatment. (A) Autophagic flux analysis of transgenic pUbi::AtC53-GFP (right panel) and pUbi::GFP-ATG8A (left panel) seedlings. **(B)** Autophagic flux analysis of endogenous AtC53 and ATG8, using AtC53 and ATG8 antibodies, respectively. **(C)** Autophagic flux analysis of transgenic pAtC53::AtC53-GFP seedlings. Col-0 or transgenic seedlings were incubated in control media or media containing 9 µM Torin1 (Tor) or 10 µg/ml tunicamycin (Tm). In addition, each treatment was supplied with 1 µm concanamycin A (conA) to visualize vacuolar degradation. Representative Western blots are displayed. Full-length and free GFP-bands from the same blot were separated due to different exposure times. In (C), * and ** correspond to short and long exposures of the same blot, respectively. Quantification of the relative intensities (Rel. Int.) of the protein bands were normalized for the total protein level of the lysate (Ponceau S). Average C53 levels and SD for n = 3 are shown. **(D-E) AtC53 flux is specifically induced upon phosphate starvation. (D)** Autophagic flux analysis of transgenic pUbi::AtC53-GFP (right panel) and pUbi::GFP-ATG8A (left panel) seedlings under carbon, nitrogen, and phosphate starvation conditions. **(E)** Autophagic flux analysis of endogenous AtC53 and ATG8, using AtC53 and ATG8 antibodies, respectively. Col-0 or transgenic seedlings were incubated in control media or media depleted with sucrose (-C), nitrogen (-N) or phosphate (-P). In addition, each treatment was supplied with 1 µm concanamycin A (conA) to visualize vacuolar degradation. Representative Western blots are displayed. Full-length and free GFP-bands from the same blot were separated due to different exposure times. Quantification of the relative intensities (Rel. Int.) of the protein bands were normalized for the total protein level of the lysate (Ponceau S). Average C53 levels and SD for n = 3 are shown. **(F) AtC53 autophagic flux is induced by various ER stress inducing conditions.** Western blot analysis of Arabidopsis transgenic seedlings expressing AtC53-GFP incubated in either control (Ctrl), sucrose-deficient medium (-C), 10 µg/ml tunicamycin (Tm), 3 h at 37°C (Heat), 2.5 µM Thapsigargin (Tg), or 50 µM Kifunensine (Kif). In addition, each treatment was supplied with 1 µm concanamycin A (conA) to visualize vacuolar degradation. (**G) HsC53 flux is induced by Torin and tunicamycin treatment.** Western blot analysis of HeLa whole cell lysates. Cells were either left untreated or treated for 16 h with 2.5 µg/ml tunicamycin (Tm) or 1.5 µM Torin (Tor) and subsequently given a recovery period of 2 h in presence or absence of 100 nM Bafilomycin A1 (BAF). C53 and BIP3 blots were run on 4-20% gradient gels and transferred to nitrocellulose membranes, LC3B blots were run on 15% gels and transferred to PVDF membranes. (* or ** indicate corresponding membranes). Quantification of the relative intensities (Rel. Int.) of the protein bands were normalized for the total protein level of the lysate (Vinculin). Average C53 levels and SD for n = 3 are shown.

### C53-ATG8 interaction is mediated by non-canonical shuffled ATG8 interacting motifs (sAIM)

Having validated C53 as an autophagy substrate, we next sought to identify its ATG8-interacting motif (AIM). For this purpose, we reconstituted the binary complex *in vitro* and determined the stoichiometry of the C53-ATG8 interaction by native mass spectrometry (nMS). Both HsC53 and AtC53 formed 1:1 and 1:2 complexes with GABARAP and ATG8A, respectively; pointing to the existence of multiple binding epitopes (Figure 3a). To map the ATG8-binding region of C53, we performed *in vitro* pull downs using truncated proteins. C53 contains an intrinsically disordered region (IDR) that bridges two a-helical domains located at the N and C termini. *In vitro* pull downs revealed that the IDR is necessary and sufficient to mediate ATG8 binding, as also confirmed with ITC and nMS experiments (Figure 3b-d, Figure S8). Multiple sequence alignment of the C53-IDR uncovered three highly conserved sites with the consensus sequence “IDWG”, representing a shuffled version of the canonical AIM sequence (W/F/Y-X-X-L/I/V) (Figure 3c, Figure S9). Mutational analysis of the three shuffled AIM sites in HsC53 and AtC53 revealed the importance of the sAIM epitopes for binding to GABARAP and ATG8, respectively; though in AtC53, an additional canonical AIM had to be mutated to fully abrogate the binding (Figure 3e). ITC experiments with the purified IDRs, as well as nMS and surface plasmon resonance (SPR) experiments with full length proteins, also supported sAIM-mediated ATG8-binding for both HsC53 and AtC53 (Figure 3f, g, Figure S10). Circular dichroism spectroscopy showed that sAIM mutants had very similar secondary structures to the wild type proteins, suggesting that lack of ATG8 binding is not due to misfolding (Figure S10c). To verify our *in vitro* results *in vivo*, we analysed the subcellular distribution of sAIM mutants in transgenic Arabidopsis lines and transfected HeLa cells. Confocal microscopy analyses showed that C53^sAIM^ mutants were not recruited into autophagosomes and had diffuse localization patterns upon ER stress induction (Figure 3h, i). Altogether these biochemical and cell biological analyses show that C53 is recruited to the autophagosomes by interacting with ATG8 via the non-canonical sAIMs.

**Figure 3.**
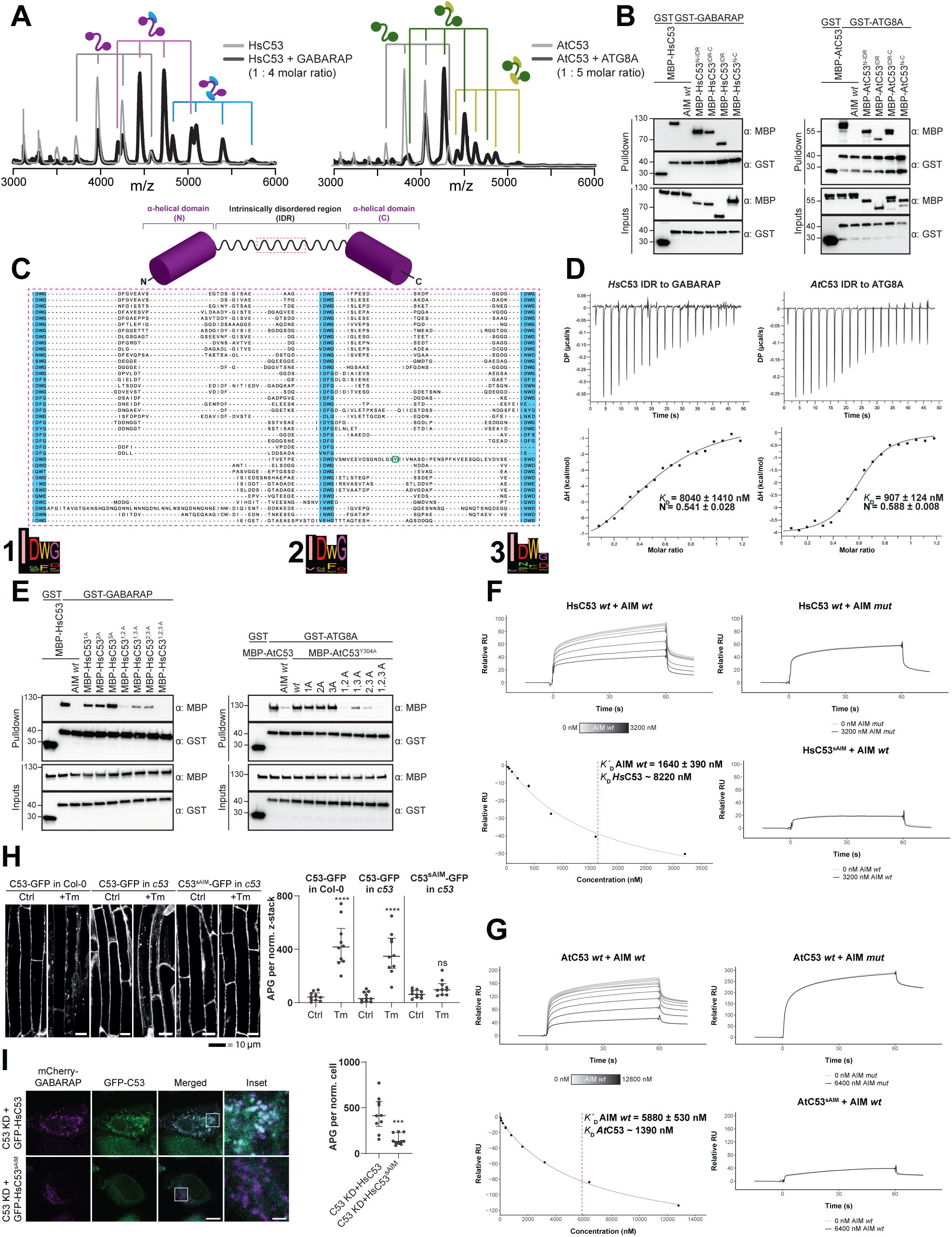
C53 interacts with ATG8 via shuffled ATG8 interacting motifs (sAIMs). (**A**) **Native mass spectrometry (nMS) analysis showing HsC53 and AtC53 form 1:1 and 1:2 complexes with GABARAP and ATG8A, respectively**. Left; nMS of HsC53 (grey) and HsC53 plus GABARAP in a 1:4 molar ratio (black). Peaks corresponding to unbound HsC53, the 1:1 complex and 1:2 complex are indicated with grey, magenta and blue, respectively. Right; nMS of AtC53 (grey) and AtC53 plus ATG8A in a 1:5 molar ratio (black). Peaks corresponding to unbound AtC53, the 1:1 complex and 1:2 complex are indicated with grey, green and yellow, respectively. Full spectra are shown in Figure S9. (**B**) *Left Panel,* HsC53 intrinsically disordered region (IDR) is necessary and sufficient to mediate the interaction with GABARAP. *Right Panel*, AtC53 IDR is necessary and sufficient to mediate the interaction with ATG8A. Bacterial lysates containing recombinant protein were mixed and pulled down with glutathione magnetic agarose beads. The AIM *wt* peptide was added at a final concentration of 200 µM. Input and bound proteins were visualized by immunoblotting with anti-GST and anti-MBP antibodies. N: N-terminal truncation; M: IDR; C: C-terminal truncation. **(C) C53 IDR has three highly conserved regions.** Protein sequence alignment of the predicted IDR amino acid sequences showed three highly conserved regions with a consensus sequence of IDWG (highlighted in blue). Y304 is highlighted in the green rectangle. The species names and the full protein sequence alignment is presented in Figure S10. (**D**) Isothermal titration calorimetry (ITC) experiments showing binding of *At*C53 and *Hs*C53 IDRs to ATG8A and GABARAP, respectively. Upper left and right panels show heat generated upon titration of *At*C53 IDR (250 µM) or *Hs*C53 IDR (250 µM) to ATG8A or GABARAP (both 40 µM). Lower left and right panels show integrated heat data (•) and the fit (solid line) to a one-set-of-sites binding model using PEAQ-ITC analysis software. Representative values of *K*_D_, N, t1H, -Tt1S, and t1G from three independent ITC experiments are reported in Table S5. (**E**) *Left Panel,* the three conserved IDWG motifs (sAIMs) in *Hs*C53 IDR mediate interaction with GABARAP. Pull downs were performed as described in (b). 1A: W269A; 2A: W294A; 3A: W312A. *Right Panel*, *At*C53 quadruple mutant cannot interact with ATG8A. In addition to the sAIM motifs, a canonical AIM (304-YEIV) also contributes to ATG8 binding. Pull downs were performed as described in (b). 1A: W276A; 2A: W287A; 3A: W335A. (**F, G**) **Surface plasmon resonance (SPR) analyses of C53-ATG8 binding.** GST-GABARAP or GST-ATG8A fusion proteins were captured on the surface of the active cell (500 RU) and GST alone was captured on the surface of the reference cell (300 RU). *Upper left Panels:* Increasing concentrations of the AIM *wt* peptide were premixed with 10 µM C53 and injected onto the sensor surface. Binding curves were obtained by subtracting the reference cell signal from the active cell signal. *lower left Panels*: Binding affinities were determined by plotting the maximum response units versus the respective concentration of the AIM *wt* peptide and the data were fitted using the Biacore T200 Evaluation software 3.1 (GE Healthcare). *Upper Right Panels:* C53 was premixed with buffer or 3600 nM of AIM *mut* peptide and injected onto the sensor surface. *lower Right Panels:* C53^sAIM^ was premixed with buffer or 3600 nM of AIM wt peptide and injected onto the sensor surface. A representative sensorgram from three independent experiments is shown. (**H**) *At*C53 quadruple mutant (sAIM) does not form puncta upon ER-stress. *left Panel*, representative confocal images of transgenic Arabidopsis seedlings expressing AtC53-GFP or AtC53^sAIM^-GFP in Col-0 wild type and *c53* mutant backgrounds. 4-day old seedlings were incubated in either control or tunicamycin (10 µg/ml) containing media. Scale bars, 10 µm. *Right Panel*, Quantification of autophagosomes (APG) per normalized Z-stacks. Bars represent the mean (± SD) of at least 10 biological replicates. (I) *Hs*C53 sAIM mutant does not form puncta upon ER-stress. *left Panel,* Confocal images of PFA fixed C53 knockdown HeLa cells transiently expressing *Hs*C53-GFP or *Hs*C53^sAIM^-GFP (green) and mCherry-GABARAP (magenta). Cells were treated for 16 h with 2.5 µg/ml tunicamycin (Tm). Scale bar, 10 µm. Inset scale bar, 2 µm. Representative images are shown. *Right Panel,* Quantification of autophagosomes (APG) per normalized cell. Bars represent the mean (± SD) of at least 10 biological replicates. Significant differences are indicated with * when p value ≤ 0.05, ** when p value ≤ 0.01, and *** when p value ≤ 0.001.

### C53 is activated by ribosome stalling during co-translational protein translocation

Next, we looked for client proteins subject to C53-mediated autophagy. Quantitative proteomics analyses of wild type and AtC53 mutant lines revealed that AtC53 mediates degradation of ER resident proteins as well as proteins passaging the ER to the cell wall, apoplast, and lipid droplets (Figure S11, Table S3, 4). These data are consistent with a recent study, showing that ER-resident proteins accumulate in a conditional mutant of mouse C53 (Yang *et al*., 2019). Since C53 is a cytosolic protein, we then explored how it senses proteotoxic stress in the ER lumen, considering four likely scenarios: C53 may collaborate with (*i*) a sensor of the unfolded protein response (UPR) (Karagoz *et al*., 2019) or (*ii*) a component of the ER-associated degradation pathway (ERAD) (Sun and Brodsky, 2019). Alternatively, it may sense clogged translocons caused by (*iii*) ribosome stalling triggered during co-translational protein translocation (Wang *et al*., 2019) or (*iv*) aberrant signal recognition particle (SRP) independent post-translational protein translocation events (Ast *et al*., 2016) (Figure S12a). To test the connection with the UPR system, we used western blot and live cell imaging based autophagic flux assays and demonstrated that AtC53 flux was already higher than wild type in Arabidopsis UPR sensor mutants *irela/b* and *bzipl7/28* (Figure S12b, c) (Kim *et al*., 2018; Koizumi *et al*., 2001). Consistently, inhibition of Ire1 activity using chemical inhibitors 4µ8c or KIRA6 also increased HsC53 puncta in HeLa cells (Figure S12d); indicating that recruitment of C53 to the autophagosomes does not depend on UPR sensors (Maly and Papa, 2014). Next, we performed colocalization analyses using model ERAD substrates. In transgenic plant lines expressing model ERAD substrates, the client proteins did not colocalize with AtC53 puncta (Figure S13a) (Shin *et al*., 2018). Likewise, the model mammalian ERAD substrates GFP-CFTRΔF508 (ERAD-C), A1AT^NHK^-GFP (ERAD-L), and INSIG1-GFP (ERAD-M) only partially colocalized with HsC53 puncta in HeLa cells (Figure S13b), suggesting C53-mediated autophagy may cross-talk with the ERAD pathway (Leto *et al*., 2019).

Next, we tested the effect of clogged translocons on C53 function. Remarkably, HsC53 significantly colocalized with the ER-targeted ribosome stalling construct ER-K20 (Wang *et al*., 2019), but not with an SRP-independent translocon clogger construct (Ast *et al*., 2016), despite both leading to clogging at the Sec61 translocon (Figure S13c). HsC53 puncta were also induced by anisomycin treatment (Figure S13d), which also leads to ribosome stalling (Wang *et al*., 2019). Consistently, silencing of HsC53 using shRNA significantly reduced lysosomal trafficking of ER-K20 (Figure S13e), suggesting C53 is activated upon ribosome stalling during co-translational protein translocation (Wamsley *et al*., 2017).

### C53 forms a tripartite receptor complex with the ufmylation E3 ligase UFL1 and its membrane adaptor DDRGK1

How is C53 recruited to the ER during ribosome stalling? Notably, C53 has been previously linked to UFL1, an E3 ligase that mediates ufmylation of stalled, ER-bound ribosomes, modifying ribosomal protein RPL26 (Walczak *et al*., 2019; Wang *et al*., 2019). To test if C53 forms a higher order receptor complex, we analysed the interaction of C53 with UFL1 and its ER membrane adaptor DDRGK1 (Gerakis *et al*., 2019). *In vitro* pull-down assays and yeast two hybrid analysis showed that AtUFL1 directly interacts with AtC53 and AtDDRGK1 (Figure 4a, Figure S14a). Consistently, AtC53 associates with DDRGK1 and UFL1 in *in vivo* coimmunoprecipitations (Figure 4b). Furthermore, co-localization of UFL1 and DDRGK1 with AtC53 in punctate structures increases upon ER stress and these puncta are delivered to the vacuole (Figure 4c, d, Figure S14b, c). Strikingly, AtC53 autophagic flux requires functional UFL1 and DDRGK1, as the number of AtC53 puncta were significantly lower in ufl1 and ddrgk1 mutants (Figure 4e, Extended Figure 14d). Altogether these data suggest C53 is recruited to the ER by forming a tripartite receptor complex with UFL1 and DDRGK1.

**Figure 4.**
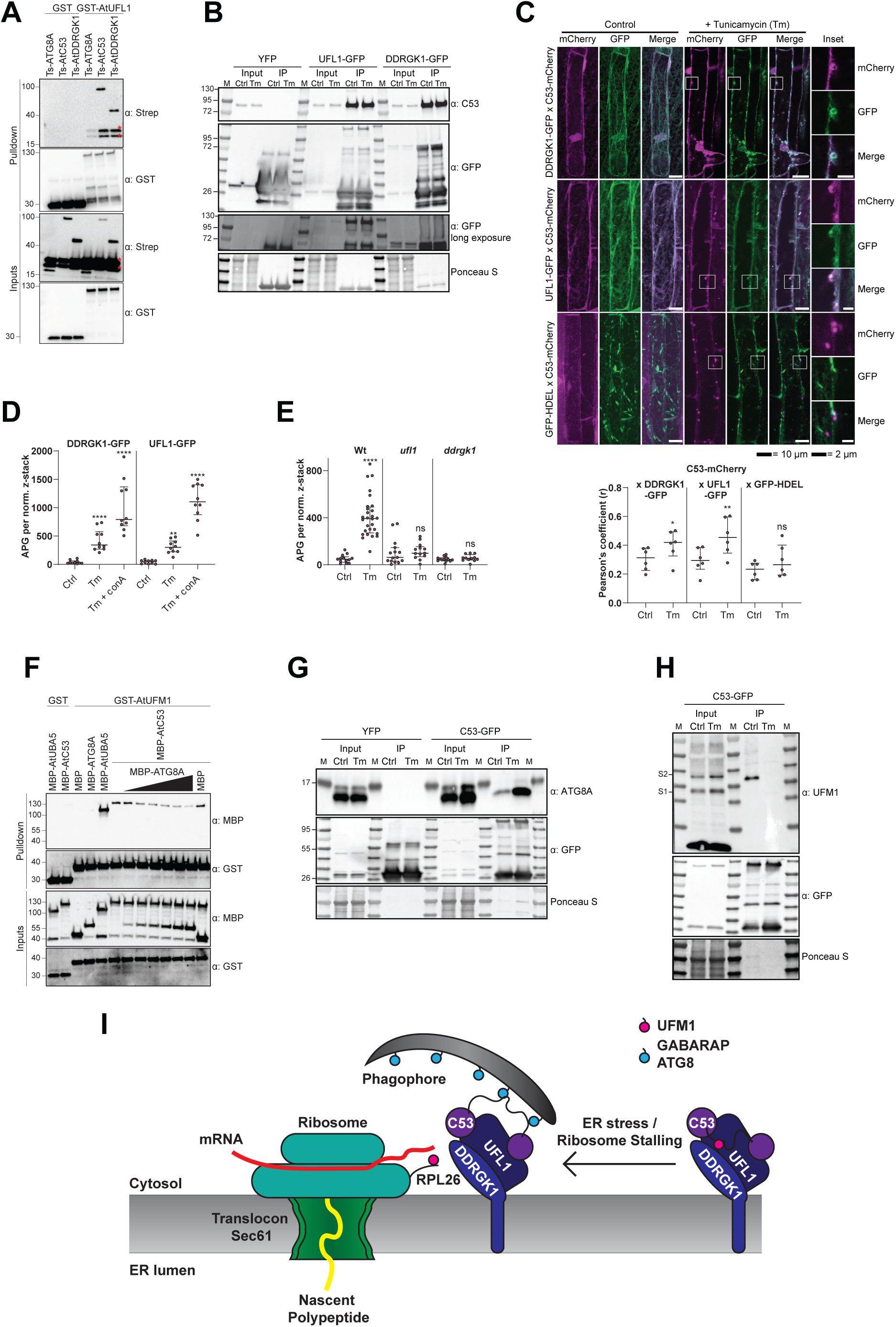
C53 forms a tripartite receptor complex and is activated by depletion of UFMt during ER stress. (A) AtUFLt interacts with AtC53 and AtDDRGKt. Bacterial lysates containing recombinant protein were mixed and pulled down with glutathione magnetic agarose beads. Input and bound proteins were visualized by immunoblotting with anti-GST and anti-Strep antibodies. Red asterisks indicate endogenous *E. coli* biotinylated proteins (B) AtC53 associates with AtUFLt and AtDDRGKt. *In vivo* co-immunoprecipitation of UFL1-GFP or DDRGK1-GFP expressing Arabidopsis seedlings incubated in either control (Ctrl) or 10 µg/ml tunicamycin (Tm) containing media. (C) AtDDRGKt and AtUFLt colocalize with AtC53 puncta upon ER stress induction. *left Upper Panel*, Co-localization analyses of confocal micrographs of wild type Col-0 roots co-expressing AtC53-mCherry (magenta) with DDRGK1-GFP, UFL1-GFP, or GFP-HDEL (green). Transgenic seedlings were incubated in either control or tunicamycin (10 µg/ml) containing media. Representative confocal images of control conditions are shown in maximum projection to emphasize ER association. Images of tunicamycin treatments are shown in single plane. Scale bars, 10 µm. Inset scale bars, 2 µm. *left lower Panel*, Pearson’s Coefficient colocalization analysis per normalized Z-scan. Bars represent the mean (± SD) of 5 biological replicates. **(D) DDRGKt and UFLt undergo vacuolar degradation upon ER stress induction.** Quantification of confocal micrographs of autophagic flux of UFL1-GFP and DDRGK1-GFP. Seedlings were incubated in either control, 10 µg/ml tunicamycin (Tm), or 10 µg/ml tunicamycin with 1 µM Concanamycin A (Tm+ConA) media. Quantification of autophagosomes (APG) per normalized Z-stacks of UFL1-GFP and DDRGK1-GFP. Bars represent the mean (± SD) of at least 10 biological replicates. **(E) AtC53 vacuolar degradation requires DDRGKt and UFLt.** Quantification of confocal images of Wildtype (Wt), *ufll*, and *ddrgkl* Arabidopsis seedlings expressing AtC53-mCherry. 6-day old seedlings were incubated in either control (Ctrl) or 10 µg/ml tunicamycin (Tm) containing media. Scale bars, 20 µm. Quantification of autophagosomes (APG) per normalized Z-stacks. Bars represent the mean (± SD) of at least 10 biological replicates. Significant differences compared to control treatment (Ctrl) are indicated with * when p value ≤ 0.05, ** when p value ≤ 0.01, and *** when p value ≤ 0.001. **(F) AtUFMt directly interacts with UFMt and this interaction becomes weaker upon increasing concentrations of ATG8A.** Bacterial lysates containing recombinant protein were mixed and pulled down with glutathione magnetic agarose beads. Input and bound proteins were visualized by immunoblotting with anti-GST and anti-MBP antibodies. The red asterisk indicates MBP-ATG8A. (G) AtC53-ATG8 association becomes stronger upon ER stress induction triggered by tunicamycin. *In vivo* co-immunoprecipitation of extracts of Arabidopsis seedlings expressing AtC53-GFP incubated in either control (Ctrl) or 10 µg/ml tunicamycin (Tm) containing media. (H) AtC53-UFMt association becomes weaker upon ER stress induction triggered by tunicamycin. *In vivo* pull downs of extracts of Arabidopsis seedlings expressing C53-GFP incubated in either control (Ctrl) or 10 µg/ml tunicamycin (Tm) containing media. Input and bound proteins were visualized by immunoblotting with the indicated antibodies. **(I) Current working model of the C53 receptor complex.** Upon ribosome stalling, UFM1 is transferred from the C53 receptor complex to the tail of RPL26, exposing the sAIMs on C53 for ATG8 binding and subsequent recruitment to the autophagosomes.

We then explored how C53 is kept inactive under normal conditions. We hypothesized that UFM1 may safeguard the C53 receptor complex under normal conditions and keep ATG8 at bay. Upon ER stress, UFM1 would be transferred to RPL26, exposing sAIMs on C53. To test this, we first tested UFM1-C53 interaction, using i*n vitro* pull downs, and showed that AtC53 can interact with AtUFM1 (Figure 4f). Furthermore, *in vitro* competition experiments suggested a competition between UFM1 and ATG8 (Figure 4f), reminiscent of the mutually exclusive UFM1 and GABARAP binding of UBA5, the E1 enzyme in the ufmylation cascade (Huber *et al*., 2019). We then performed *in vivo* co-immunoprecipitation experiments during ER stress. Consistent with our hypothesis and *in vitro* data, ER stress led to depletion of UFM1 and enhanced AtC53-ATG8 interaction (Figure 4g, h, Figure S14e). Altogether, these data suggest that the two ubiquitin-like proteins UFM1 and ATG8 compete with each other for association with the C53 receptor complex (Figure 4i).

### C53 is crucial for ER stress tolerance

Finally, we examined if C53 is physiologically important for ER stress tolerance. First, we tested if C53 plays a general role in autophagy using carbon and nitrogen starvation assays. Carbon and nitrogen starvation are typically used to characterize defects in bulk autophagy responses (Marshall and Vierstra, 2018). In contrast to the core autophagy mutants *atg5* and *atg2*, CRISPR-generated *Atc53* mutants did not show any phenotype under carbon or nitrogen starvation conditions (Figure 5a, b). However, consistent with increased flux, *Atc53* mutants were highly sensitive to phosphate starvation, which has been shown to trigger an ER stress response (Naumann *et al*., 2018) (Figure 5c, Figure S15a). Similarly, in both root length and survival assays, *Atc53* mutants were sensitive to tunicamycin treatment (Figure 5d, Figure S15b, c). In addition, ufmylation machinery mutants (Figure 5e), including *ufll* and *ddrgkl*, were also sensitive to tunicamycin treatment but insensitive to carbon and nitrogen starvation (Figure 5f, Figure S15d, e). Lastly, the *Marchantia polymorpha c53* mutant was also sensitive to tunicamycin, suggesting C53 function is conserved across the plant kingdom (Figure S15f). We then performed complementation assays using wild type AtC53 and the AtC53^sAIM^ mutant. AtC53 expressing lines behaved like wild type plants in tunicamycin supplemented plates (Figure 5g). However, AtC53^sAIM^ mutant did not complement the tunicamycin sensitivity phenotype, and had significantly shorter roots (Figure 5g, Figure S15g). Parallel to analysing C53-mediated ER homeostasis in plants, stress tolerance assays in HeLa cells showed that silencing of HsC53 led to an induction of Bip3 chaperone protein levels (Figure S15h), indicating increased ER stress. Complementation of *Hsc53* silenced lines with HsC53-GFP dampened Bip3 expression (Figure S15h). Altogether, these results demonstrate that C53 coordinated ER-phagy is crucial for ER stress tolerance in plant and mammalian cells.

**Figure 5.**
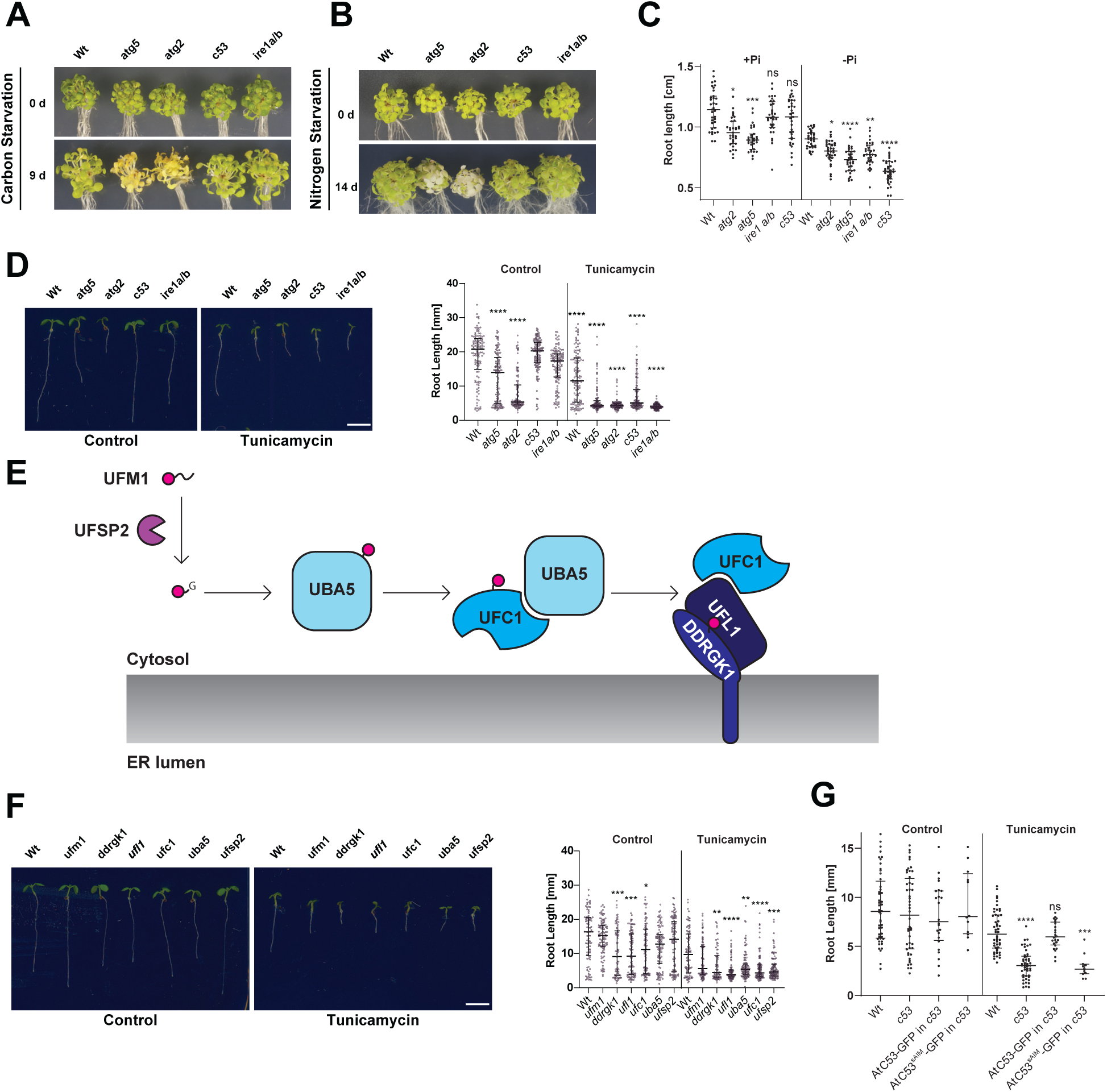
C53 is crucial for ER stress tolerance. (**A**) ***Atc53* mutant is insensitive to carbon starvation.** Phenotypes before (0 d) and after 9 days carbon starvation (9 d) of seven-day-old seedlings, n 2’ 20 seedlings per genotype. (**B**) ***Atc53* mutant is insensitive to nitrogen starvation.** Phenotypes before (0 d) and after 14 days nitrogen starvation (14 d) of seven-day-old seedlings, n 2’ 20 seedlings per genotype. **(C) *Atc53* mutant is sensitive to phosphate starvation.** Root-length quantification of seven-day-old seedlings which were transferred to media with or without Pi supplement (+Pi, -Pi), and imaged after 2 days. (**D**) **Atc53 mutants are sensitive to ER stress induced by tunicamycin**. Root-length quantification of 7-day old seedlings grown on half strength MS media without sucrose treated with 100 ng/mL tunicamycin (Tm). Bottom Panel, Root-length quantification of 7-day old seedlings. n;c125 seedlings per genotype and treatment. *Le/t Panel*, Example of 7-day old seedlings grown in described conditions. Scale bars = 5 mm. Left, non-treated seedlings. Right, seedlings grown at 100 ng/mL Tm. *Right Panel,* Root length of each genotype was compared pairwise with the wildtype (Col-0) for each specific treatment condition. **(E) Main molecular players in the ufmylation pathway.** UFSP2: UFM1 specific protease 2 that matures UFM1, exposing the terminal glycine residue. UBA5: the E1 activating enzyme, UFC1: E2 conjugating enzyme, UFL1: E3 ligase (**F**) **Ufmylation pathway mutants are sensitive to ER stress triggered by tunicamycin.** Root length quantification of 7-day old seedlings grown on half strength MS media without sucrose treated with 100 ng/mL tunicamycin (Tm). *Le/t panel,* Root length quantification of 7-day old seedlings. n;c100 seedlings per genotype and treatment. *Right Panel,* Representative images of 7-day old seedlings grown in described conditions. Scale bars, 5 mm. To the left are non-treated seedlings, to the right are seedlings grown at 100 ng/mL Tm. **(G) AtC53^sAIM^ mutant does not complement tunicamycin sensitivity phenotype.** Root length quantification of indicated 7-d old seedlings grown on half strength MS media without sucrose in control conditions (Ctrl) or treated with 100 ng/mL tunicamycin (Tm). T1 transgenic lines were used. n=12 seedlings per genotype and treatment. Data represent the median with its interquartile range. Root length of each genotype was compared pairwise with the wildtype (Col-0) for each treatment condition. Significant differences compared to control treatment (Ctrl) are indicated with * when p value ≤ 0.05, ** when p value ≤ 0.01, and *** when p value ≤ 0.001.

## Discussion

In conclusion, our data show that C53 forms an ancient autophagy receptor complex that is closely connected to the ER quality control system via the ufmylation pathway. During ER stress, UFM1 is depleted from the C53-receptor complex, exposing the sAIM motifs on C53 for ATG8 binding. Notably, the Sec61 translocon and ER-associated ribosomes are not targeted by C53, pointing to further factors complementing the C53/UFM1/DDRDPK system of selective ER-phagy.

Our findings highlight C53 mediated ER-phagy is a central player operating at the interface of key quality control pathways, controlling ER homeostasis across different kingdoms of life. Consistently, recent genome wide CRISPR screens identified ufmylation as a major regulator of ER-phagy, the ERAD pathway, and viral infection (Kulsuptrakul *et al*., 2019; Leto *et al*., 2019; Liang *et al*., 2020). Excitingly, using fluorescent reporter lines and genome wide CRISPRi screens, Liang *et al*., showed that ufmylation plays a major role in regulating starvation induced ER-phagy. They showed that both DDRGK1 and UFL1 are critical for starvation induced ER-phagy; whereas C53 mutants did not show any ER-phagy defects (Liang *et al*., 2020). Our results using stable transgenic organisms show that C53 mediated autophagy is not activated by carbon or nitrogen starvation that are typically used to activate bulk autophagy. C53 is activated by phosphate starvation, but this is not due starvation but because of the ER stress triggered during phosphate starvation. Furthermore, C53 and the ufmylation machinery is asymptomatic during carbon or nitrogen starvation. Additionally, Liang *et al*., show ufmylation works together with known ER-phagy receptors such as Fam134B. Whether C53 also works with other ER-phagy receptors needs further investigation. Differences in our and Liang et. al.’s findings may imply cell-type specific differences and highlight the need for further studies to resolve the discrepancies.

C53 and ufmylation are essential for mammalian development (Gerakis *et al*., 2019). Defects in C53 receptor complex have been associated with various diseases including liver cancer, pancreatitis, and cardiomyopathy (Gerakis *et al*., 2019). Our results suggest C53 and ufmylation is also critical for stress tolerance in plants, but they are not essential for development; suggesting plants have evolved compensatory mechanisms during adaptation to sessile life. Future comparative studies could reveal these mechanisms and help us develop sustainable strategies for promoting ER proteostasis during stress in mammals and plants.

## Acknowledgements

We thank R. Strasser, M. Schuldiner, R. Kopito, J. Christianson, K. Mukhtar, S. Howell, D. Hofius, R. Vierstra, Y. Ye, W. Yarbrough and T. Ueda for sharing plasmids or plant lines. We acknowledge the Vienna BioCenter Core Facilities GmbH (VBCF) facilities Plant Sciences (J. Jez), Electron Microscopy (T. Heuser, S. Jakob, M. Brandstetter), and Protein Technologies (P. Stolt-Bergner, A. Sedivy, J. Neuhold, A. Lehner) as well as the GMI/IMBA/IMP Protein Chemistry Core facility (M. Madalinski, E. Roitinger, K. Stejskal, R. Imre) and A. Schleiffer for their help with the experiments. The SPR and ITC equipment were kindly provided by the EQ-BOKU VIBT GmbH and the BOKU Core Facility Biomolecular & Cellular Analysis. We thank members of the *Vienna Biocenter Ubiquitin Club* for fruitful discussions. This work has been funded by the Vienna Science and Technology Fund (WWTF) through project LS17-O47 (YD, TC), the Austrian Science Fund (FWF): P32355 (YD), P3O4O1-B21 (SM), I3O33-B22 (AD), Unidocs Fellowship (AS), ERC grant No.646653 (S.M.), and the Austrian Academy of Sciences. M.M. is financially supported by The Financial Supports for Young Scientists (WULS-SGGW) International Research Scholarship Fund No. BWM 315/2O18. We thank Claudine Kraft, Elif Karagoz, James Watson and Youssef Belkhadir for critical evaluation of the manuscript. We also acknowledge Sabbi Lall, Life Science Editors for editing assistance.

## Author Contributions

MS, LP, TC and YD conceived and designed the project. MS, ES, AM, MC, CN, MM and CL, performed plant related experiments. AG, AA, AS performed or contributed to human cell culture related experiments. RB performed native mass spectrometry experiments. KC performed the yeast two hybrid assays. LP, VM, ET and HC performed *in vitro* biochemical and biophysical assays. YZ, BL performed electron microscopy experiments. GD, MS performed quantitative mass spectrometry experiments. AD, IS, SA, LJ, KM, FI, SM, TC, YD supervised the project. MS, LP, AG and YD wrote the manuscript with input from all the authors.

---

**Figure S1.**
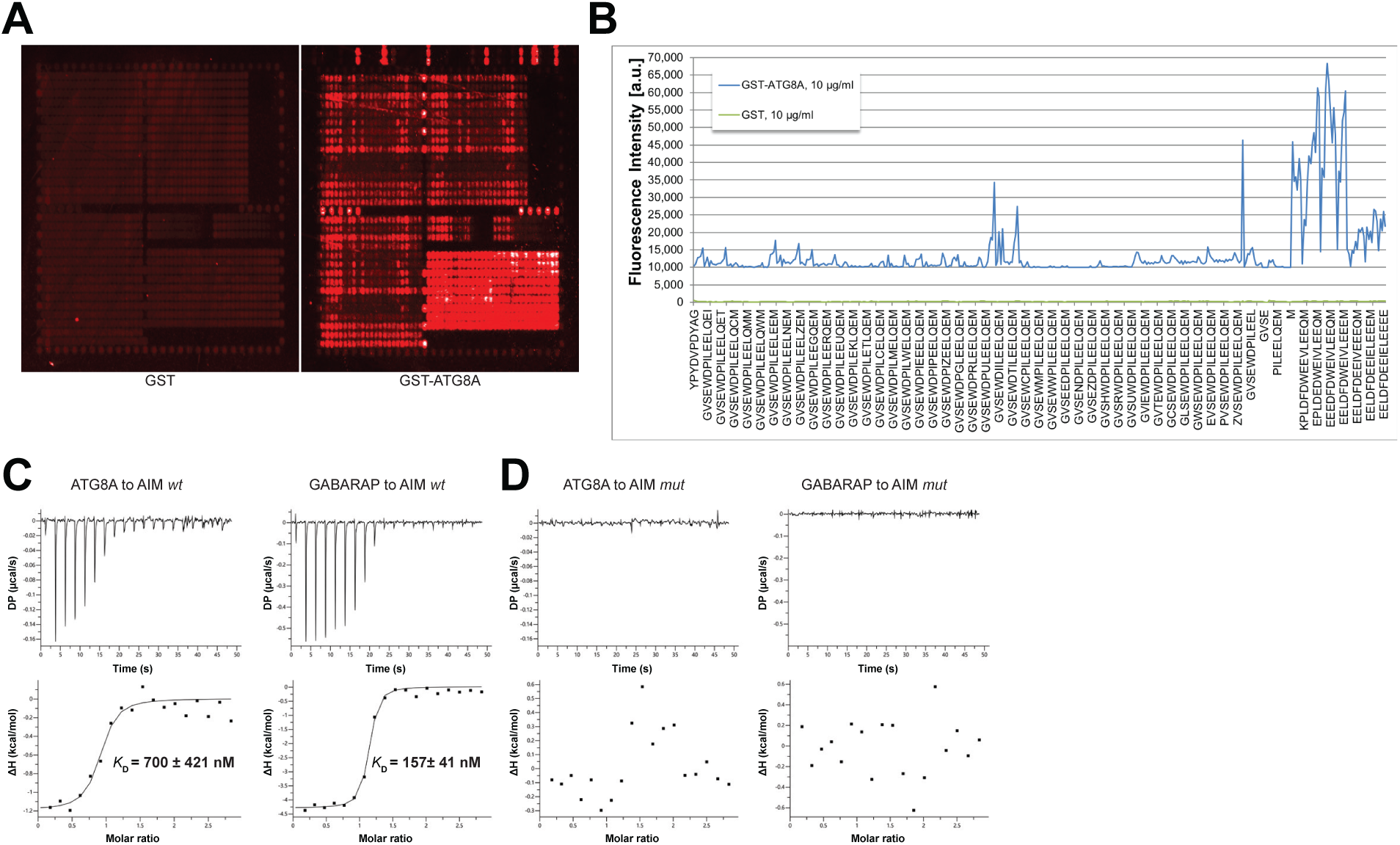
Identification of high affinity AIM peptides for peptide competition coupled immunoprecipitation mass spectrometry and *in vitro* pull-down experiments. (a) Qualitative analysis of peptide array results. A library of ATG8-interacting motif peptides (See Table S1), were spotted onto an array and incubated with GST or GST-ATG8A. **(b) Quantification of peptide array results for selected AIM peptides.** The AIM peptide (AIM *wt*) (EEEDFDWEIVLEEEM) showed the highest fluorescence intensity. (c) Isothermal titration calorimetry (ITC) experiments showing binding of AIM *wt* to ATG8A and GABARAP. Upper left and right panels show heat generated upon titration of AIM *wt* (600 µM) to ATG8A or GABARAP (both 40 µM). Lower left and right panels show integrated heat data (•) and the fit (solid line) to a one-set-of-sites binding model using PEAQ-ITC analysis software. Representative values of *K*_D_, N, ΔH, -TΔS and ΔG from three independent ITC experiments are reported in Table S5. (d) Isothermal titration calorimetry (ITC) experiments showing that the AIM mutant peptide (AIM *mut*) (EEEDFDAEIALEEEM) cannot bind to ATG8A or GABARAP. Upper left and right panels show heat generated upon titration of *AlM mut* (600 µM) to ATG8A or GABARAP (both 40 µM). Lower left and right panels show integrated heat data (•) from PEAQ-ITC analysis software.

**Figure S2.**
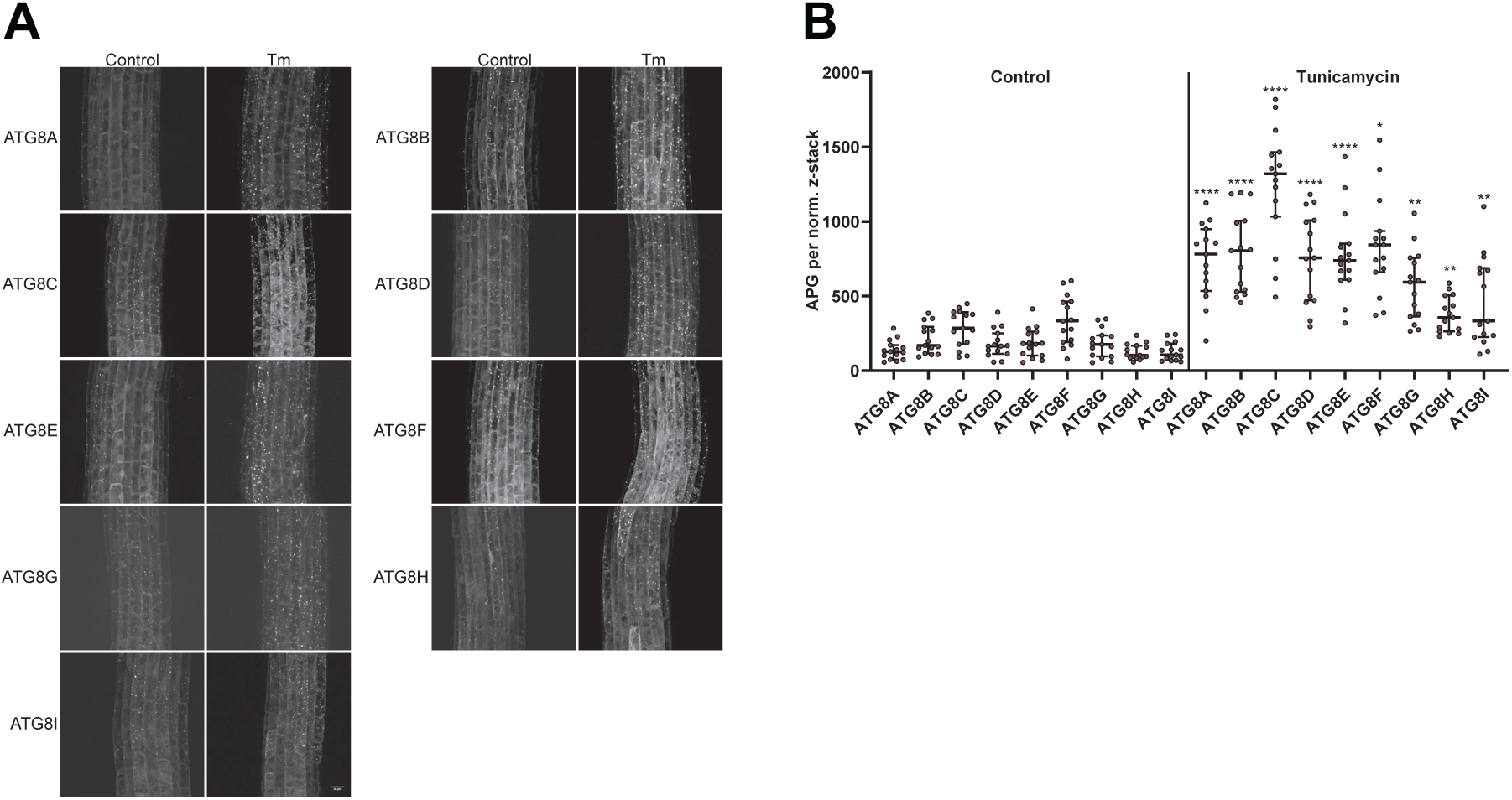
All Arabidopsis ATG8 isoforms are induced by tunicamycin-triggered ER stress. (a) Representative confocal images of transgenic Arabidopsis seedlings expressing GFP-ATG8A to I in Col-0 background. 6-day old seedlings were incubated in either control or tunicamycin (10 µg/ml) containing media. **(b)** Quantification of the autophagosomes (APG) per normalized Z-stack. Bars represent the mean (± SD) of at least 10 biological replicates. Significant differences between control and tunicamycin-treated samples are indicated with * when p value ≤ 0.05, ** when p value ≤ 0.01, and *** when p value ≤ 0.001.

**Figure S3.**
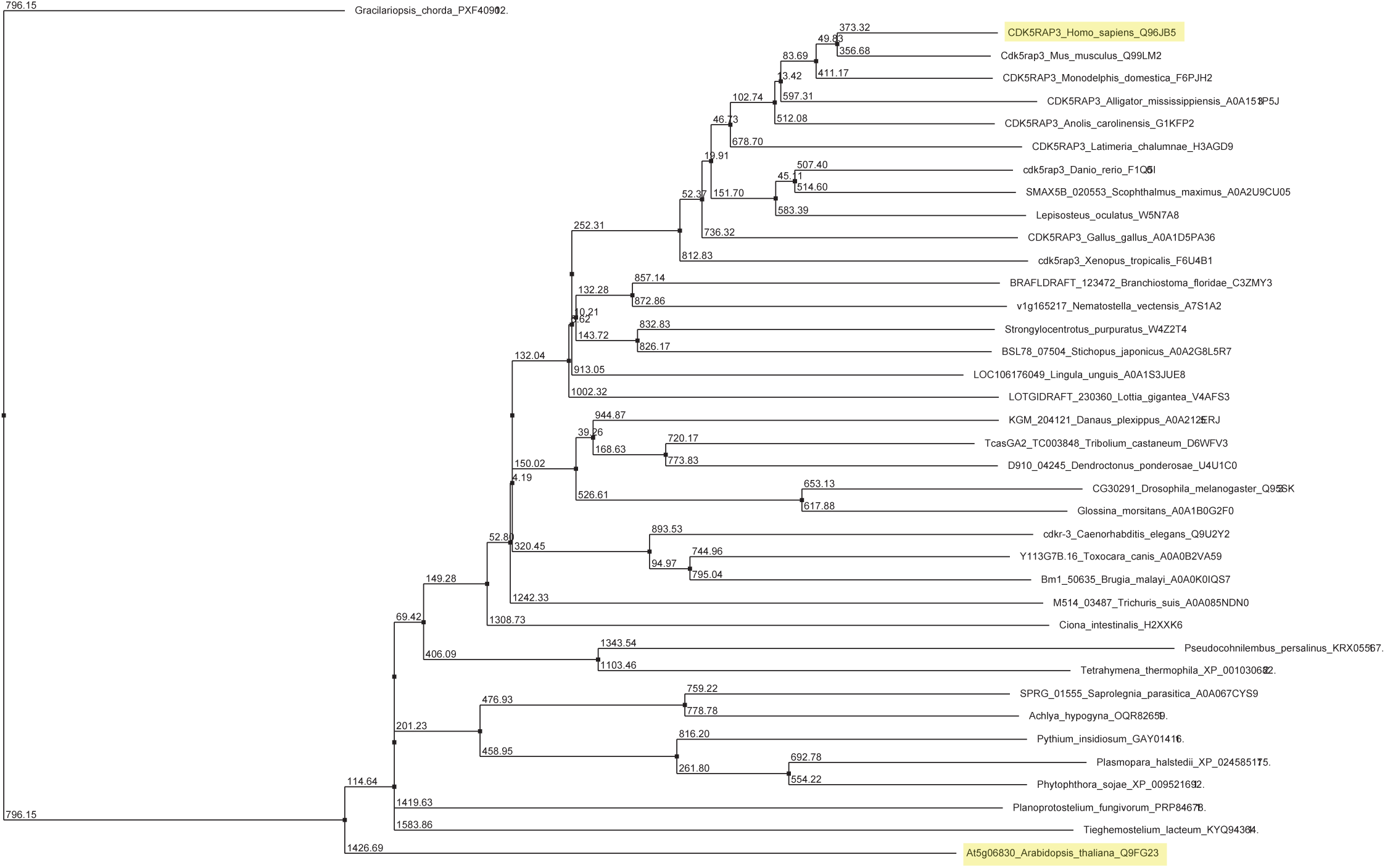
Unrooted maximum likelihood phylogenetic tree of C53 homologs. The tree was generated from protein sequence alignments of C53 homologs from 38 species. The bootstrap support for each node is indicated. The tree was calculated with Jalview 2.11.0 from distance matrices determined from aggregate BLOSUM 62 scores using Average Distance (UPGMA) algorithm.

**Figure S4.**
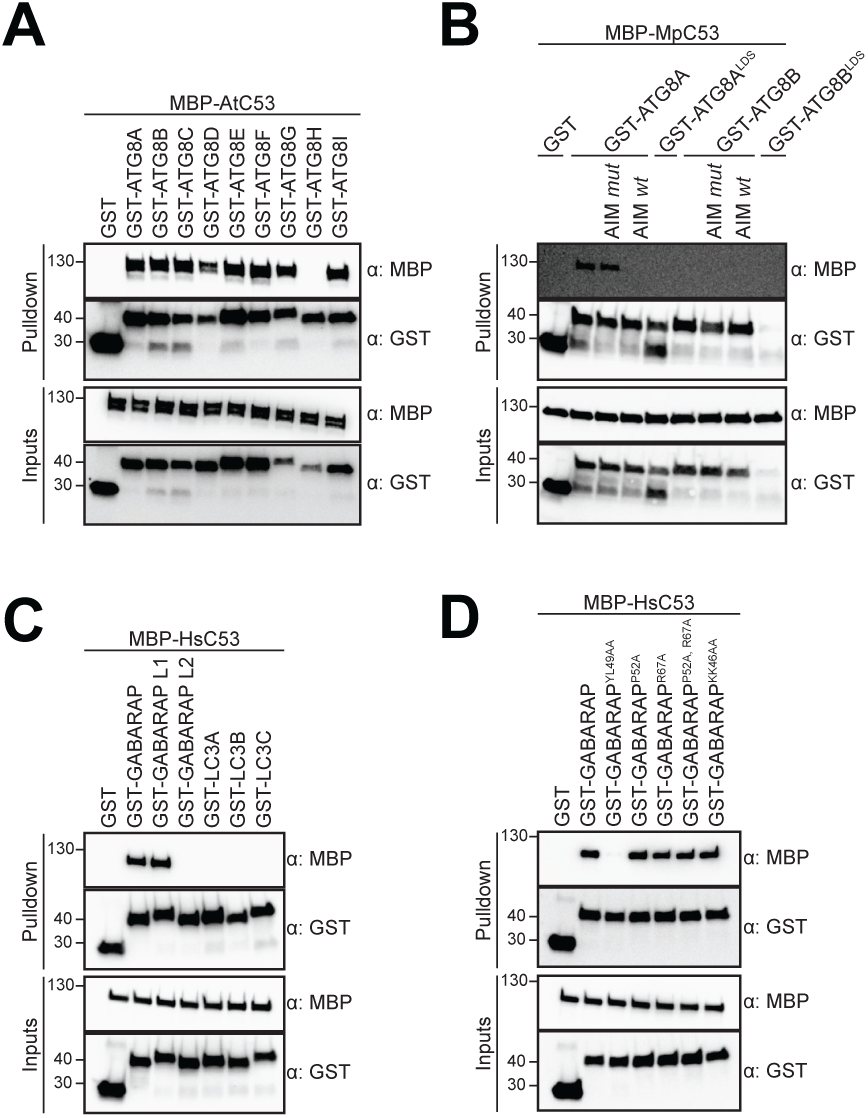
C53 binds ATG8 isoforms from plants and humans in an AIM-dependent manner. (**a**) AtC53 interacts with AtATG8 in an isoform specific manner. */n vitro* pull down with all ATG8 isoforms of *Arabidopsis thaliana* (At) shows that AtC53 can interact with eight out of nine ATG8 isoforms. Bacterial lysates containing recombinant protein were mixed and pulled down with glutathione magnetic agarose beads. (b) MpC53 interacts with MpATG8 isoforms in a specific manner. */n vitro* pull down with both ATG8 isoforms of *Marchantia polymorpha* (Mp) shows that MpC53 can interact with one out of two ATG8 isoforms. Bacterial lysates containing recombinant protein were mixed and pulled down with glutathione magnetic agarose beads. Before MpC53 pull down, peptides AIM *wt* and AIM *mut* were added to a final concentration of 200 µM. Input and bound proteins were visualized by immunoblotting with anti-GST and anti-MBP antibodies. (**c**) **HsC53 interacts with GABARAP and GABARAP Ll.** Bacterial lysates containing recombinant protein were mixed and pulled down with glutathione magnetic agarose beads. Input and bound proteins were visualized by immunoblotting with anti-GST and anti-MBP antibodies. **(d) HsC53 interacts with GABARAP via the LIR Docking Site (LDS).** Mutating the W site to a YL49AA mutation (Marshall *et al*., 2019) prevents binding of GABARAP to C53. However, mutating the L position to P52A or R67A (Marshall *et al*., 2019), or mutating KK64AA (which mediates the interaction with the atypical LIR motif found in UBA5 (Huber *et al*., 2019)) did not prevent C53 binding. Bacterial lysates containing recombinant protein were mixed and pulled down with glutathione magnetic agarose beads. Input and bound proteins were visualized by immunoblotting with anti-GST and anti-MBP antibodies.

**Figure S5.**
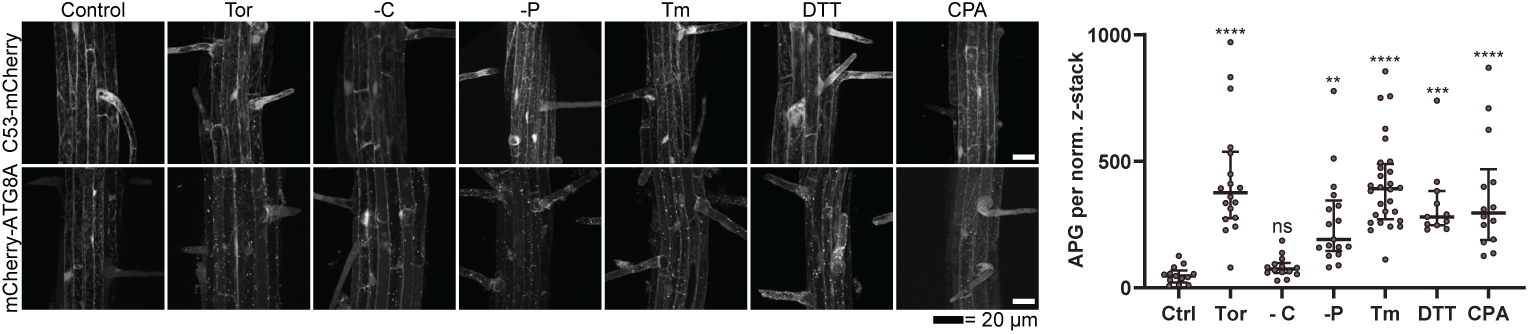
Analysis of AtC53 puncta under various stress conditions revealed induction of C53 puncta upon ER stress. *Left Panel*, representative confocal images of transgenic Arabidopsis seedlings expressing C53-mCherry and mCherry-ATG8A in Col-0 background. 6-day old seedlings were incubated in either control (Ctrl), sucrose deficient (-C), or phosphate deficient (-P) media, or media containing 9 µm Torin1 (Tor), 10 µg/ml tunicamycin (Tm), 15 µM CPA, or 2.5 mM OTT. The TOR kinase inhibitor Torin and ER stress inducing conditions -P, OTT, Tm, and CPA triggered formation of both C53 and ATG8 puncta, whereas carbon starvation only triggered ATG8 puncta. Scale bars, 20 µm. *Right Panel*, Quantification of autophagosomes (APG) per normalized Z-stack. Bars represent the mean (± SO) of at least 10 biological replicates. Significant differences compared to control treatment (Ctrl) are indicated with * when p value ≤ 0.05, ** when p value ≤ 0.01, and *** when p value ≤ 0.00. ns=not significant.

**Figure S6.**
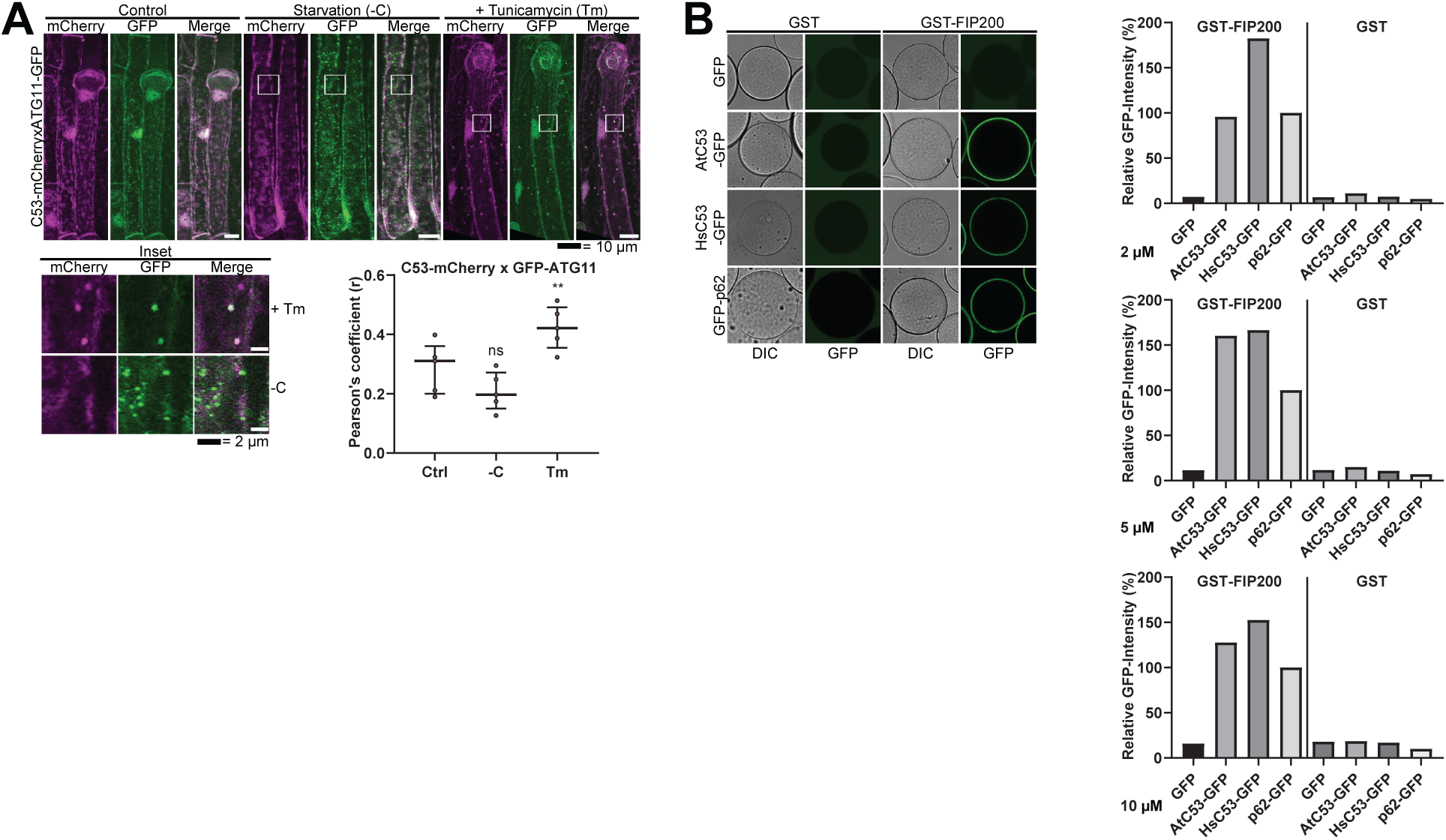
C53 binds selective autophagy adaptor ATG11. **(a) C53 puncta partially colocalize with ATG11-GFP during ER stress**. *Upper Panel*, Co-localization analyses of single plane confocal images obtained from transgenic Arabidopsis roots co-expressing C53-mCherry (magenta) with ATG11-GFP (green). 4-day old seedlings were incubated in either control, sucrose deficient (-C), or 10 µg/ml tunicamycin (Tm) containing media. Scale bars, 20 µm. *Lower Left Panel,* Inset Scale bar, 2 µm. *Lower Right Panel*, Pearson’s Coefficient (r) analysis of the colocalization of C53-mCherry with ATG11-GFP. C53 only colocalized with ATG11 under ER stress treatments. Bars represent the mean (± SD) of at least 5 biological replicates. Significant differences are indicated with * when p value = 0.05, ** when p value = 0.01, and *** when p value = 0.001. ns= not significant **(b) AtC53 and HsC53 directly interact with mammalian ATG11 homolog FIP200.** GSH beads were coated with GST or GST-FIP200. Excess GST or GST-FIP200 was washed off, and beads were incubated with 2, 5 or 10 µM of recombinant GFP (negative control), GFP-p62 (positive control), AtC53-GFP, or HsC53-GFP. Beads were equilibrated for 1 hour and imaged using a confocal microscope. *Right Panel*, Quantification of C53 binding to FIP200. The graphs on the right show the average GFP intensity of beads n 2’ 15, normalized to the signal of GFP-p62.

**Figure S7.**
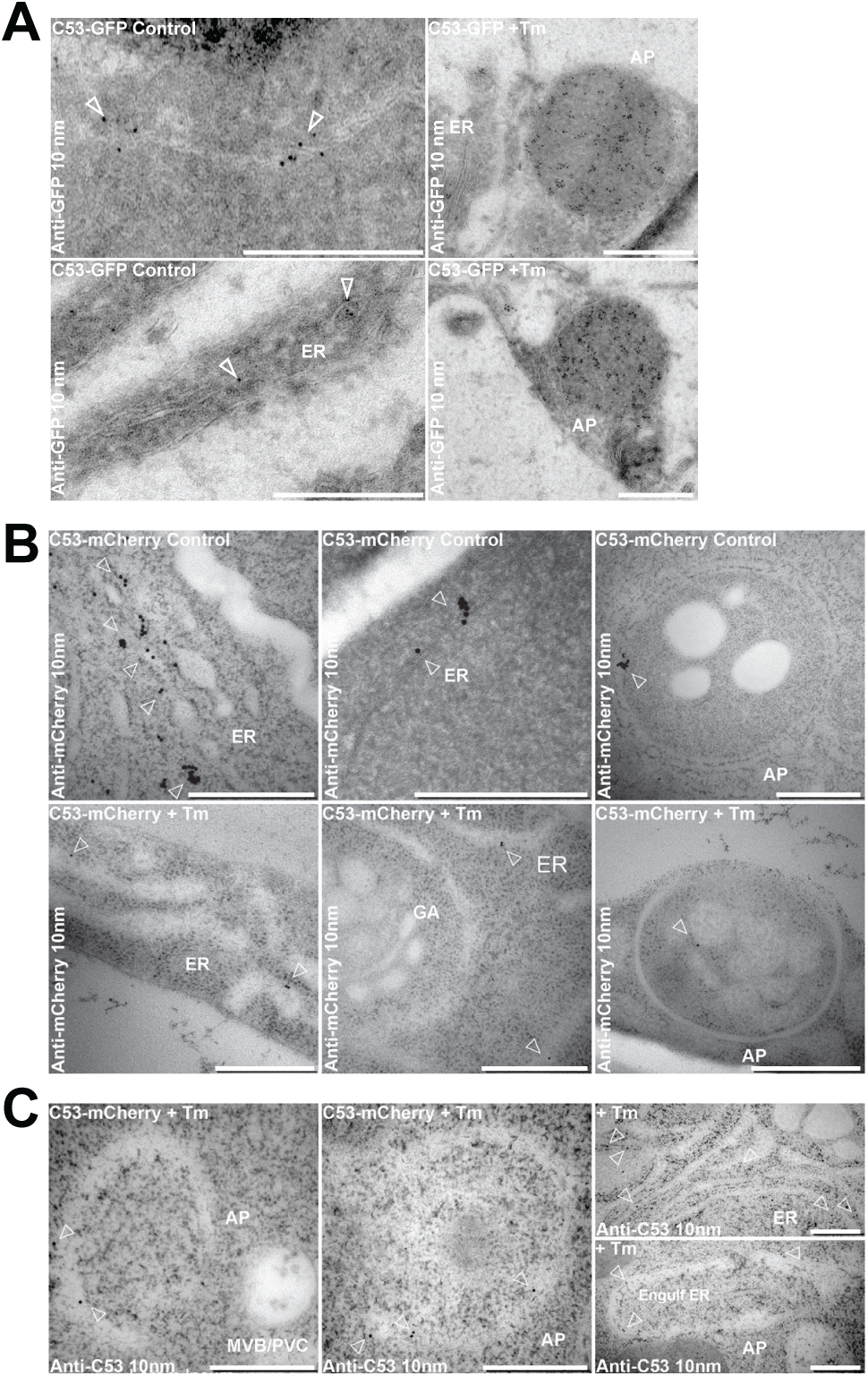
Electron micrographs showing that C53 localizes to the ER and autophagosomes during ER stress. (a-c) lmmunogold labelling of high-pressure frozen, 5-day old *Arabidopsis* roots treated with (Tm) or without (control) 10 µg/ml tunicamycin for 6 hours. Arrowheads indicate 10 nm gold particles. Scale bars, 500 nm. **(a)** C53-GFP transgenic roots were prepared for immunogold electron microscopy analysis using GFP antibody. **(b)** C53-mCherry transgenic roots were prepared for immunogold electron microscopy analysis using RFP antibody. **(C)** Col-0 non-transgenic roots were prepared for immunogold electron microscopy analysis using C53-antibody. ER= Endoplasmic reticulum, AP=autophagosome, MVB/PVC=multivesicular body

**Figure S8.**
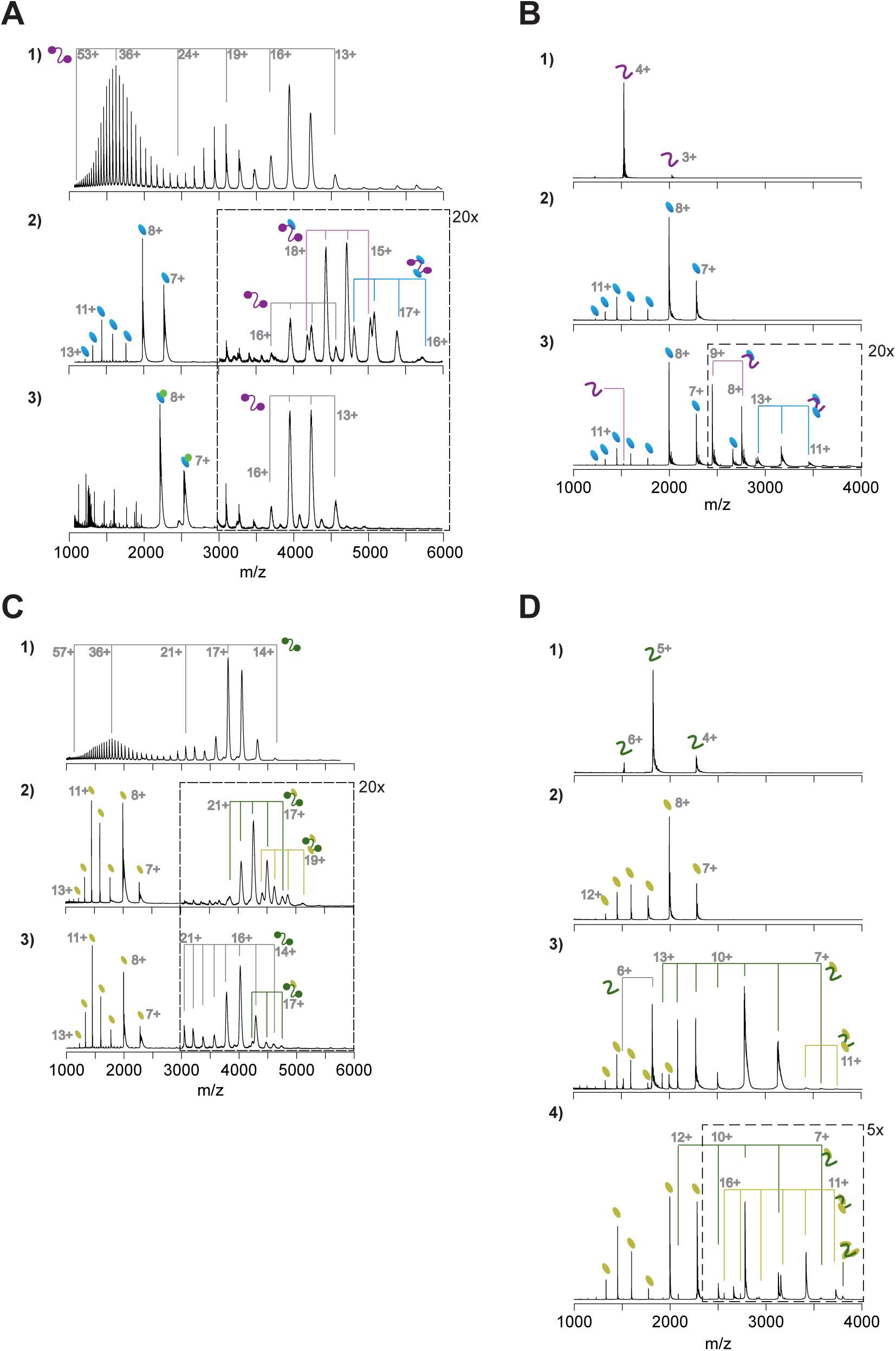
Native mass spectrometry analyses of HsC53-GABARAP and AtC53-ATG8A interactions. (a) Native mass spectrometry (nMS) of **(1)** HsC53 (5 µM), **(2)** HsC53 (5 µM) + GABARAP (20 µM), and **(3)** HsC53 (5 µM) + GABARAP (10 µM) + AIM *wt* peptide (10 µM). **(1)** HsC53 is a monomeric protein, which presents in charge states 13+ to 53+. The trimodal shape of the charge state distribution is diagnostic of a protein that exists in compact and extended conformations (Beveridge *et al*., 2014). **(2)** Binding of HsC53 to GABAPRAP is observed in 1:0, 1:1 and 1:2 ratios. **(3)** All GABARAP is now present in a 1:1 complex with the AIM peptide, which prevents it from binding to HsC53. HsC53 is present in the unbound state. **(b)** nMS of **(1)** HsC53-IDR (2.5 µM), **(2)** GABARAP (5 µM), and **(3)** HsC53-IDR (2.5 µM) + GABARAP (12.5 µM). Binding of HsC53-IDR to GABARAP is observed in 1:1 and 1:2 ratios. **(c)** nMS of **(1)** AtC53 (1 µM), **(2)** AtC53 (1 µM) + AtG8A (5 µM), and **(3)** AtC53^sAIM^ (1 µM) + AtG8A (5 µM). **(1)** AtC53 is a monomeric protein, which presents in charge states 14+ to 57+. The trimodal shape of the charge state distribution is diagnostic of a protein that exists in compact and extended conformations (Beveridge *et al*., 2014). **(2)** Binding of AtC53 to AtG8A is observed in 1:1 and 1:2 ratios. **(3)** AtC53^sAIM^ is mainly present in the unbound state, with small amounts of 1:1 complex formed with AtG8A. **(d)** nMS of **(1)** AtC53-IDR (2.5 µM), **(2)** AtG8A (10 µM), **(3)** AtC53-IDR (2.5 µM) + AtG8A (10 µM), and **(4)** AtC53-IDR (2.5 µM) + AtG8A (20 µM). Binding of AtC53-IDR to ATG8A is observed in 1:1 and 1:2 ratios, with a small amount of 1:3 binding at 1:8 molar excess of ATG8A.

**Figure S9.**
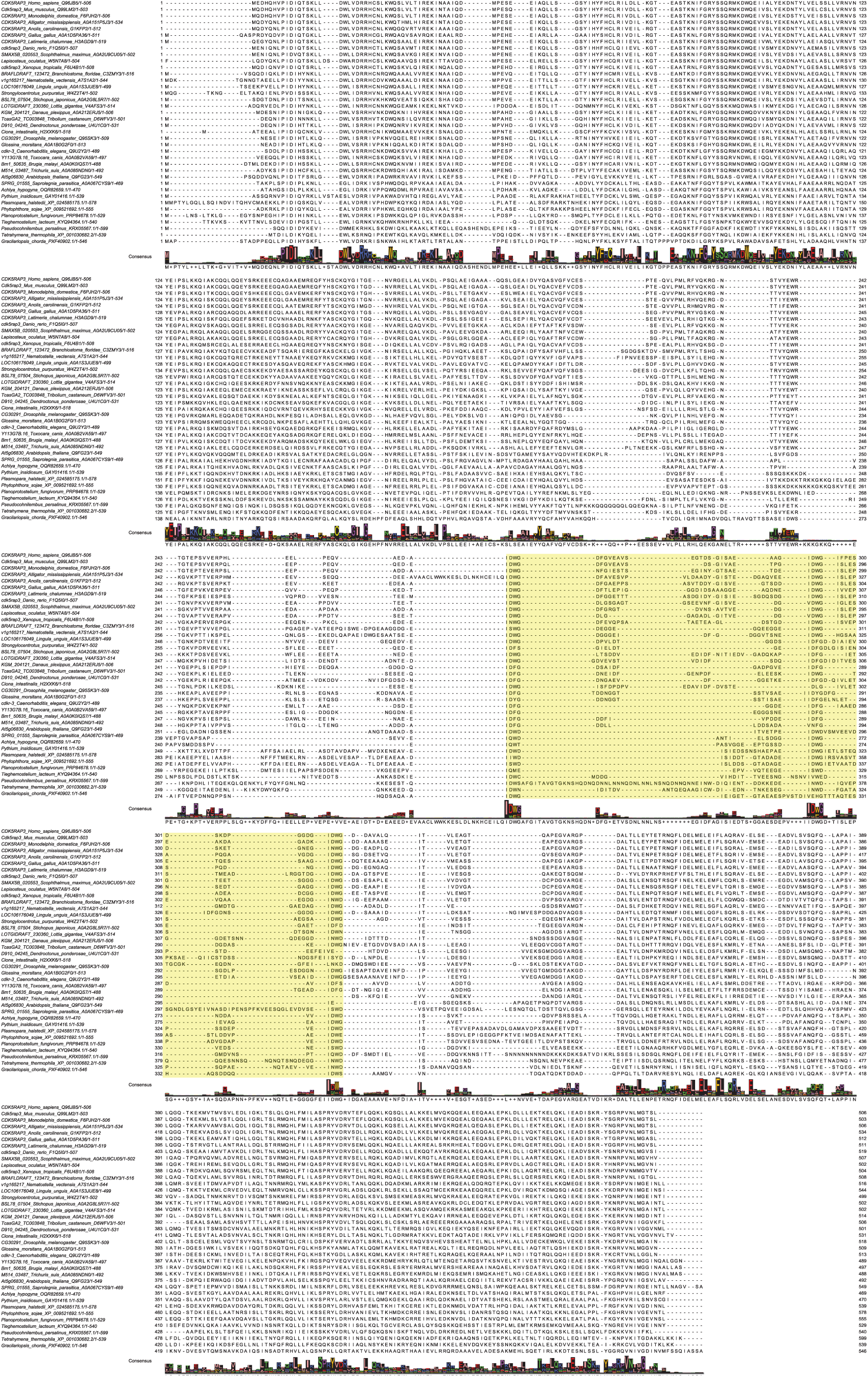
Multiple sequence alignment of C53 homologs. Protein sequences of C53 homologs from selected multicellular eukaryotic species were aligned using Clustal Omega server (Madeira *et al*., 2019) and processed with Jalview. Residue numbers are labelled according to the HsC53 sequence. The multiple sequence alignment used in Figure 3D is highlighted in yellow.

**Figure S10.**
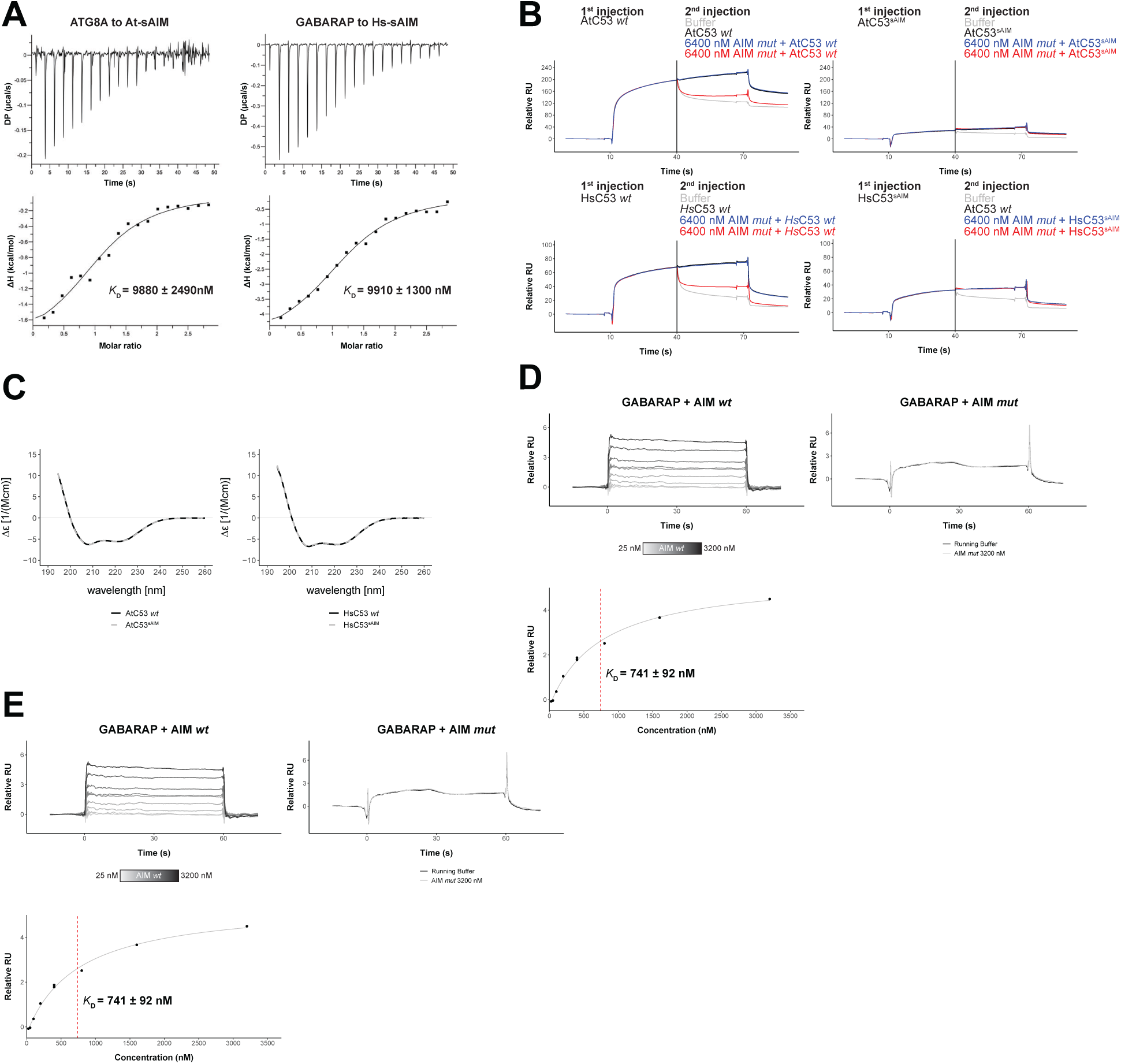
Biophysical characterization of sAIM mediated C53-ATG8 interaction. (a) Isothermal titration calorimetry (ITC) experiments showing binding of At-sAIM and Hs-sAIM peptides to ATG8A and GABARAP, respectively. Upper left and right panels show heat generated upon titration of At-sAIM (EPLDFDIDWDLEEEM) (600 µM) or Hs-sAIMC53 (EPLDFDIDWGLEEEM) (600 µM) to ATG8A or GABARAP (both 40 µM). Lower left and right panels show integrated heat data (•) and the fit (solid line) to a one-set-of-sites binding model using PEAQ-ITC analysis software. Representative values of *K*_D_, N, 11H, -T11S and 11G from three independent ITC experiments are reported in Table S5. **(b)** AtC53 and HsC53 sAIM mutants do not interact with ATG8A or GABARAP, respectively. *Upper Panels,* GST-ATG8A or *Lower Panels,* GST-GABARAP were captured on the surface of the active cell (500 RU) and GST was captured on the surface of the reference cell (300 RU). The 2 flow cells were exposed to the ligands *Upper left* AtC53, *Upper right* AtC53^sAIM^, *Lower Left*, HsC53 and *Lower right,* HsC53^sAIM^ with 4 sets of double consecutive injections (1^st^ set: 10 µM ligand, running buffer; 2^nd^ set: 10 µM ligand, 10 µM ligand; 3^rd^ set: 10 µM ligand, 10 µM ligand + 6.4 µM AIM *wt* peptide; 4^th^ set: 10 µM protein, 10 µM protein + 6.4 µM AIM *mut* peptide. Binding curves were obtained by subtracting the reference cell from the active cell. A representative sensorgram from two independent experiments is shown. **(c) AtC53^sAIM^ and HsC53^sAIM^ mutants have similar secondary structure compared to AtC53 and HsC53, respectively.** *Left Panel*, Far-UV circular dichroism (CD) spectra of AtC53 *wt* (black line) and its variant AtC53^sAIM^ (grey dashed line). *Right Panel*, Far-UV circular dichroism (CD) spectra of HsC53 *wt* (black line) and its variant HsC53^sAIM^ (grey dashed line). **(D) Quantification of the binding affinity of AIM *wt* to GABARAP.** GST-GABARAP fusion protein was captured on the surface of the active cell (500 RU) and GST was captured on the surface of the reference cell (300 RU). Representative sensorgrams from at least 3 independent experiments are shown. *Upper Left Panel*, Multi-cycle kinetics experiment with increasing concentrations of the AIM *wt* peptide. Binding curves were obtained by double referencing (i.e. reference cell signal subtracted from active cell signal; subtraction of buffer injection). 400 nM* = Internal replicate. *Upper Right Panel*, Injection of running buffer (grey) and 3600 nM AIM *mut* peptide (black). *Lower Left Panel*, Plot of steady-state response units versus the respective concentration of the AIM *wt* peptide, which was used to determine the dissociation constant *K*_D_. **(E) Quantification of the binding affinity of AIM *wt* to ATG8A.** GST-ATG8A fusion protein was captured on the surface of the active cell (500 RU) and GST was captured on the surface of the reference cell (300 RU). Representative sensorgrams from at least 3 independent measurements are shown. *Upper Left Panel*, Multi-cycle kinetics experiment with increasing concentrations of the AIM *wt* peptide. Binding curves were obtained by double referencing (i.e. reference cell signal subtracted from active cell signal; subtraction of buffer injection). 400 nM* = Internal replicate. *Upper right Panel*, Injection of running buffer (grey) and 3600 nM AIM *mut* peptide (black). *Lower Left Panel*, Plot of steady-state response units versus the respective concentration of the AIM *wt* peptide which was used to determine the dissociation constant *K*_D_.

**Figure S11.**
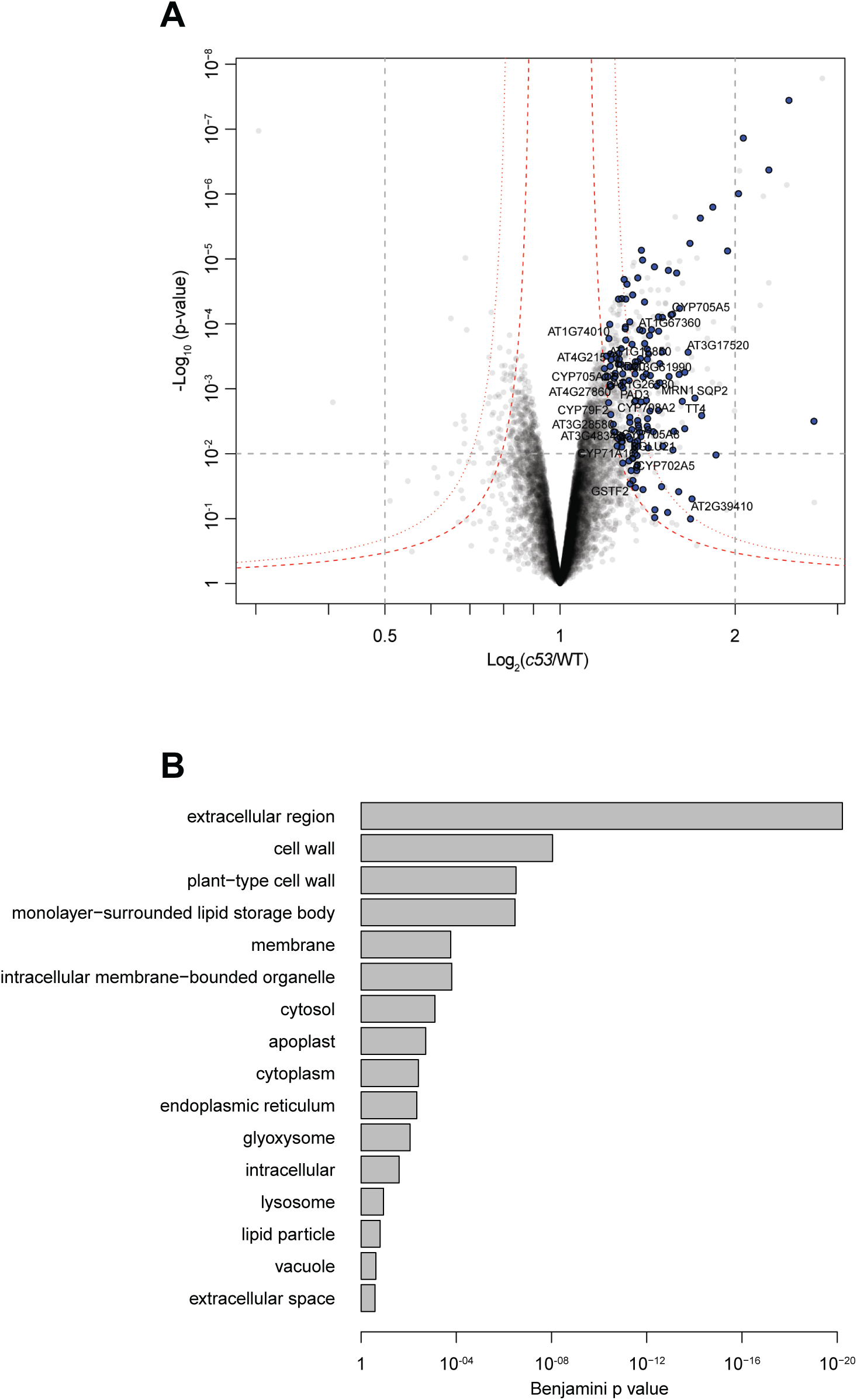
Quantitative proteomics analyses of AtC53 mediated degradation. (a) Volcano plot showing proteins that are accumulating in *Atc53 mutants*. Names of ER resident proteins are shown. Proteins that are labelled with blue either reside or mature at the ER. **(b) GO analysis of proteins accumulating in *Atc53*.** See Table S3 and Table S4 for details.

**Figure S12.**
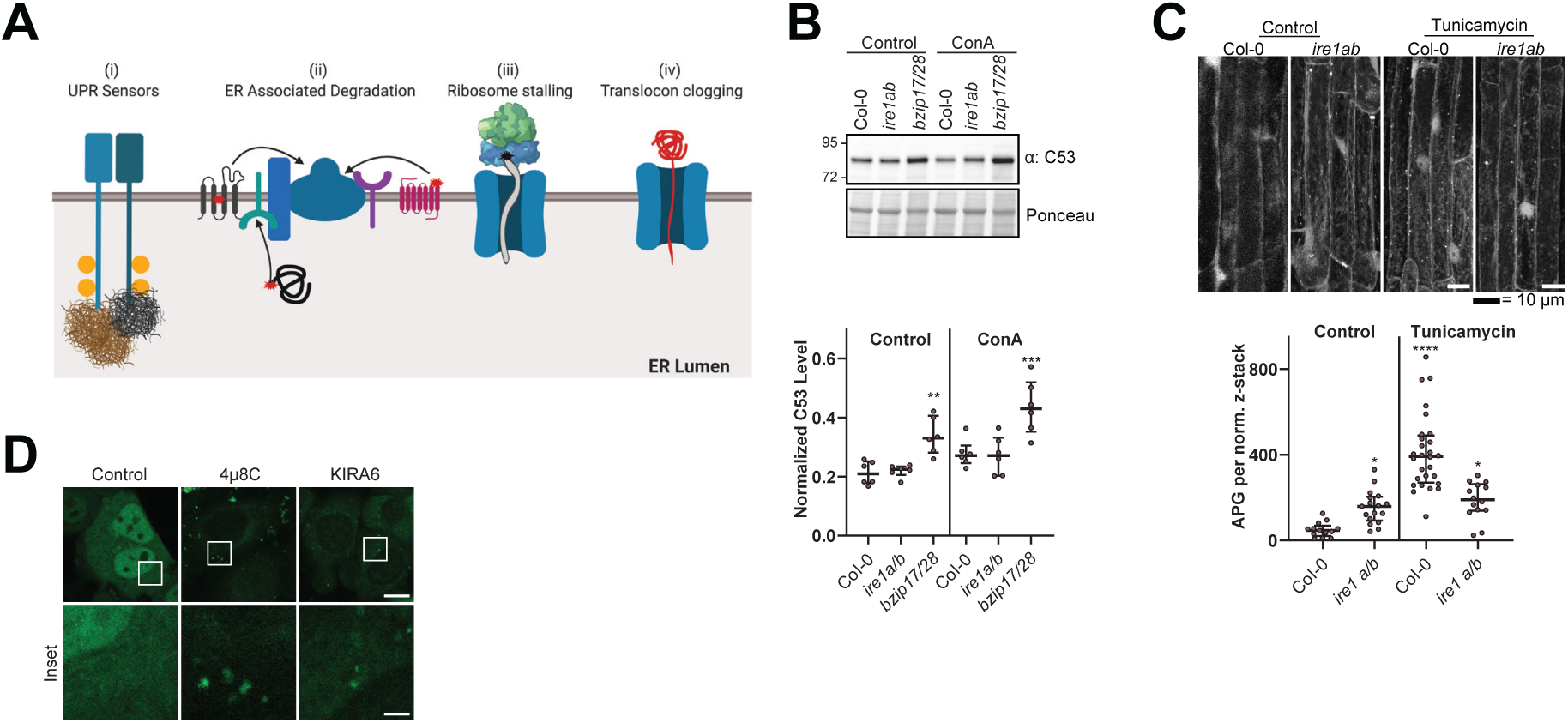
C53 is not activated by associating with UPR sensors. **(a) Cartoon explaining the four scenarios we tested to understand the mechanism of activation of C53. (b) AtC53 flux is enhanced in Arabidopsis UPR sensor mutants**. *Upper Panel*, representative western blot image of autophagic flux analysis of C53 in Col-0 wild type, *irela/b*, and *bzipl7/28* double mutants. Seedlings were incubated in either control (Ctrl) or 1 µM concanamycin A (ConA) containing medium for 16 h. Proteins extracted from whole seedlings were analysed by immunoblotting with anti-C53 antibody. Total proteins were analysed by Ponceau S staining. *Lower Panel*, Quantification of the intensities of the C53 bands normalized to the total protein level of the lysate (Ponceau S). Average C53 levels and SD for n = 6 are shown. Significant differences are indicated with * when p value ≤ 0.05, ** when p value ≤ 0.01, and *** when p value ≤ 0.001. **(c) AtC53 puncta are already induced in *irela/b* without ER stress.** *Upper Panel,* Confocal micrographs of autophagic flux of transgenic seedlings expressing C53-mCherry in Col-0 wild type and the *irela/b* double mutant. Seedlings were incubated in either control media or media containing 10 µg/ml tunicamycin (Tm). Representative confocal images in single plane are shown. Scale bars, 10 µm. *Lower Panel,* Quantification of autophagosomes (APG) per normalized Z-stack. Bars represent the mean (± SD) of at least 10 biological replicates. **(d) Chemical inhibition of IRE1 activity enhances HsC53 autophagic flux in HeLa cells.** Confocal micrographs of PFA fixed HeLa cells transiently expressing HsC53-GFP. Cells were either left untreated (Control) or treated for 1 h with 5µM 4µ8C (IRE1 RNase activity inhibitor) or 1µM KIRA6 (IRE1 kinase activity inhibitor). Inhibitor treatments led to the depletion of HsC53 from the nucleus and puncta formation. Scale Bar, 20 µm. Representative images are shown.

**Figure S13.**
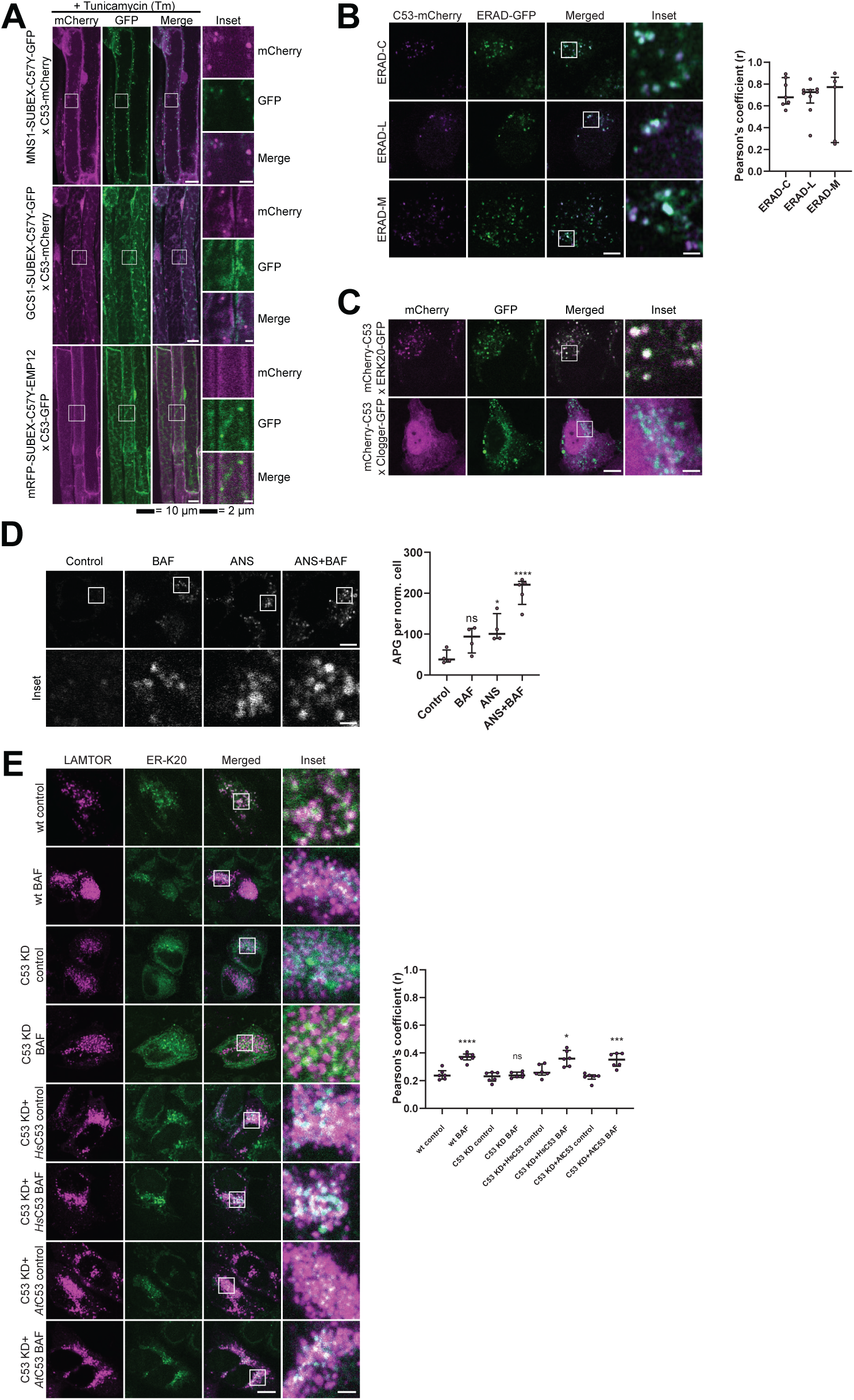
C53 is activated upon ribosome stalling during co-translational protein translocation. (a) AtC53 does not co-localize with ERAD substrates. *Left Upper Panel*, Colocalization analysis from confocal micrographs of C53-mCherry or C53-GFP (magenta/green) transgenic lines crossed with MNS1-SUBEX-C57Y-GFP (green), GCS1-SUBEX-C57Y-GFP (green), or mRFP-SUBEX-C57Y-EMP12 (magenta). Transgenic seedlings were incubated in either control or tunicamycin (10 µg/ml) containing media. Representative confocal images of control conditions are shown as maximum projection images to demonstrate the ER localization of the substrates. Images of tunicamycin treatments are shown in single plane. Scale bar, 10 µm. Inset scale bar, 2 µm. **(b) HsC53 partially co-localizes with ERAD substrates.** *Left Panel,* Confocal images of PFA fixed HeLa cells transiently co-expressing C53-mCherry with GFP-CFTRΔF508 (ERAD-C), A1AT^NHK^-GFP (ERAD-L), and INSIG1-GFP (ERAD-M). Scale Bar, 20µm. *Right Panel,* Pearson’s Coefficient analysis of the co-localization. (**c) HsC53 colocalize with ribosome stalling construct ER-K20.** Confocal images of PFA fixed HeLa cells co-expressing ER-K20-GFP or the clogger construct with mCherry-HsC53. (d) HsC53 puncta are induced by anisomycin treatment. *Left Panel,* Confocal images of PFA fixed HeLa cells transiently expressing C53-GFP. Cells were either left untreated or treated for 15 min with 200 nM anisomycin (ANS) and subsequently given a recovery period of 2 h in the presence or absence of 100 nM bafilomycin A1 (BAF). Scale Bar, 20 µm. *Right Panel,* Quantification of C53 puncta. (e) HsC53 mediates degradation of the ribosome stalling construct ER-K20. *Upper Panel,* Confocal images of PFA fixed HeLa cells transiently expressing ER_K20-GFP (green) and miRFP-LAMTOR1 (magenta). HsC53 knock down (KD) cells were complemented with HsC53 or AtC53. Cells were either left untreated or treated for 2 h with 100 nM bafilomycin A1 (BAF). Scale Bar, 20 µm. Representative images are shown. *Lower Panel:* Quantification of the colocalization of Lamtor1 with ER-K20 in knock down and complemented cells.

**Figure S14.**
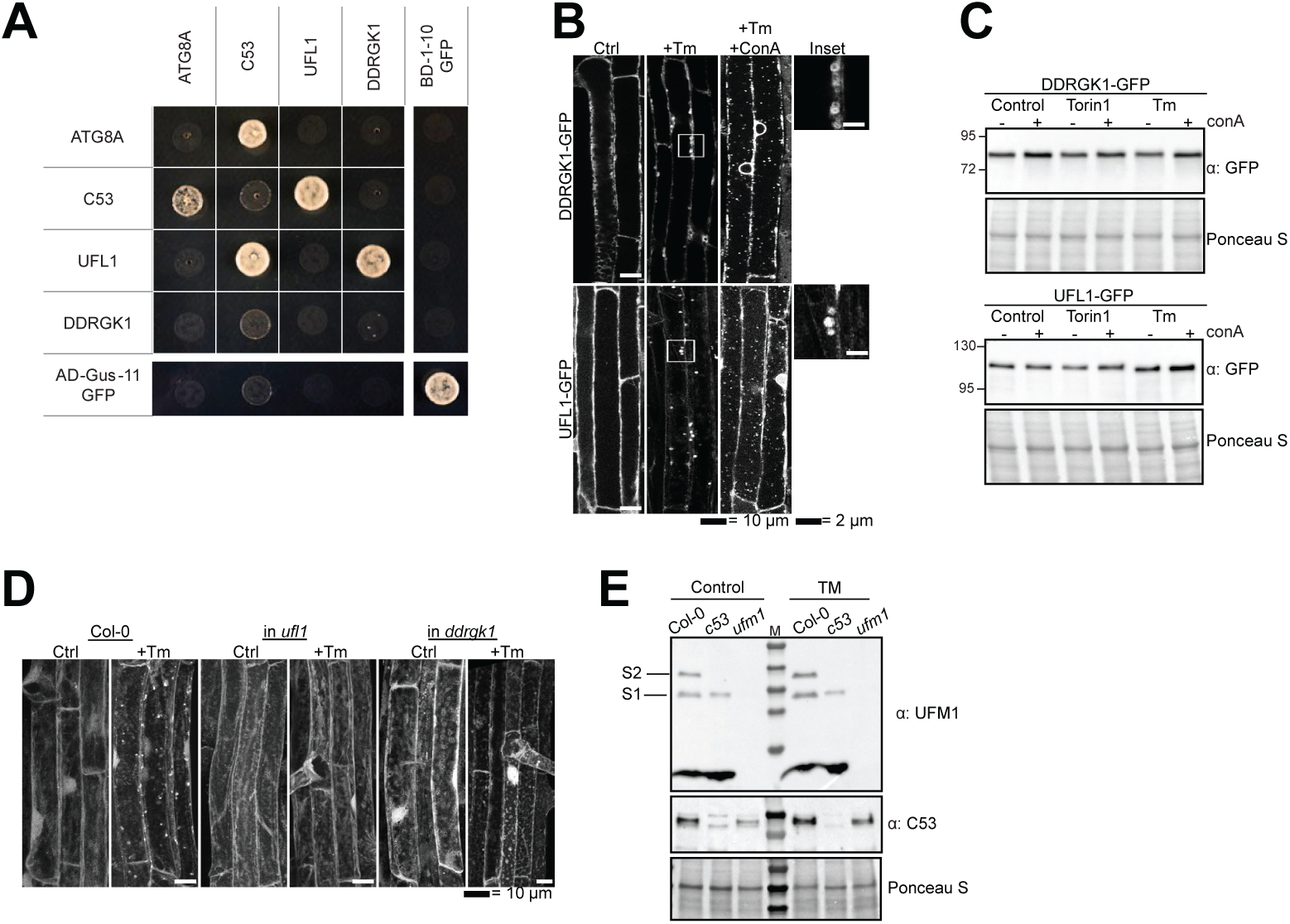
C53, DDRGKt, and UFLt form a tripartite receptor complex. **(a) pairwise Yeast two hybrid analysis of Arabidopsis thaliana UFLt, C53, DDRGKt, and ATG8A.** Combinations of pGADT7 (prey vector) and pGBKT7 (bait vector) vectors carrying the indicated genes were transformed in yeast. After mating, yeast was selected for growth on (SD)-Leu/-Trp, (SD)-Leu/-Trp/-His and (SD)-Leu/- Trp/-His/-Ade plates (displayed). Split GFP was used as a positive control. **(b, c) DDRGKt and UFLt undergo vacuolar degradation upon ER stress. (b)** Confocal micrographs of transgenic seeds expressing UFL1-GFP and DDRGK1-GFP. Seedlings were incubated in either control, 10 µg/ml tunicamycin (Tm) or 10 µg/ml tunicamycin and 1 µM concanamycin A (Tm+ConA). Representative confocal images in single plane are shown. Scale bars, 10 µm. Inset scale bars, 2 µm. **(c)** Representative western blots of UFL1-GFP and DDRGK1-GFP upon ER stress treatments. Seedlings were incubated in either control, 10 µg/ml tunicamycin (Tm) or 10 µg/ml tunicamycin with 1 µM concanamycin A (Tm+ConA). Protein extracts from whole seedlings were analysed by immunoblotting with anti-GFP. Total proteins were analysed by Ponceau S staining. **(d) AtC53 puncta formation requires UFLt and DDRGKt.** Representative confocal images of Col-0, *ufll*, and *ddrgkl* Arabidopsis seedlings. 6-day old seedlings were incubated in either control (Ctrl) or 10 µg/ml tunicamycin (Tm) containing media. In contrast to wild type, AtC53 did not form puncta in DDRGK1 and UFL1 mutants. For quantification, see Figure 3f. **(e) Validation of UFMt antibody used in 3g.** S1 and S2 correspond to ufmylated RPL26 bands.

**Figure S15.**
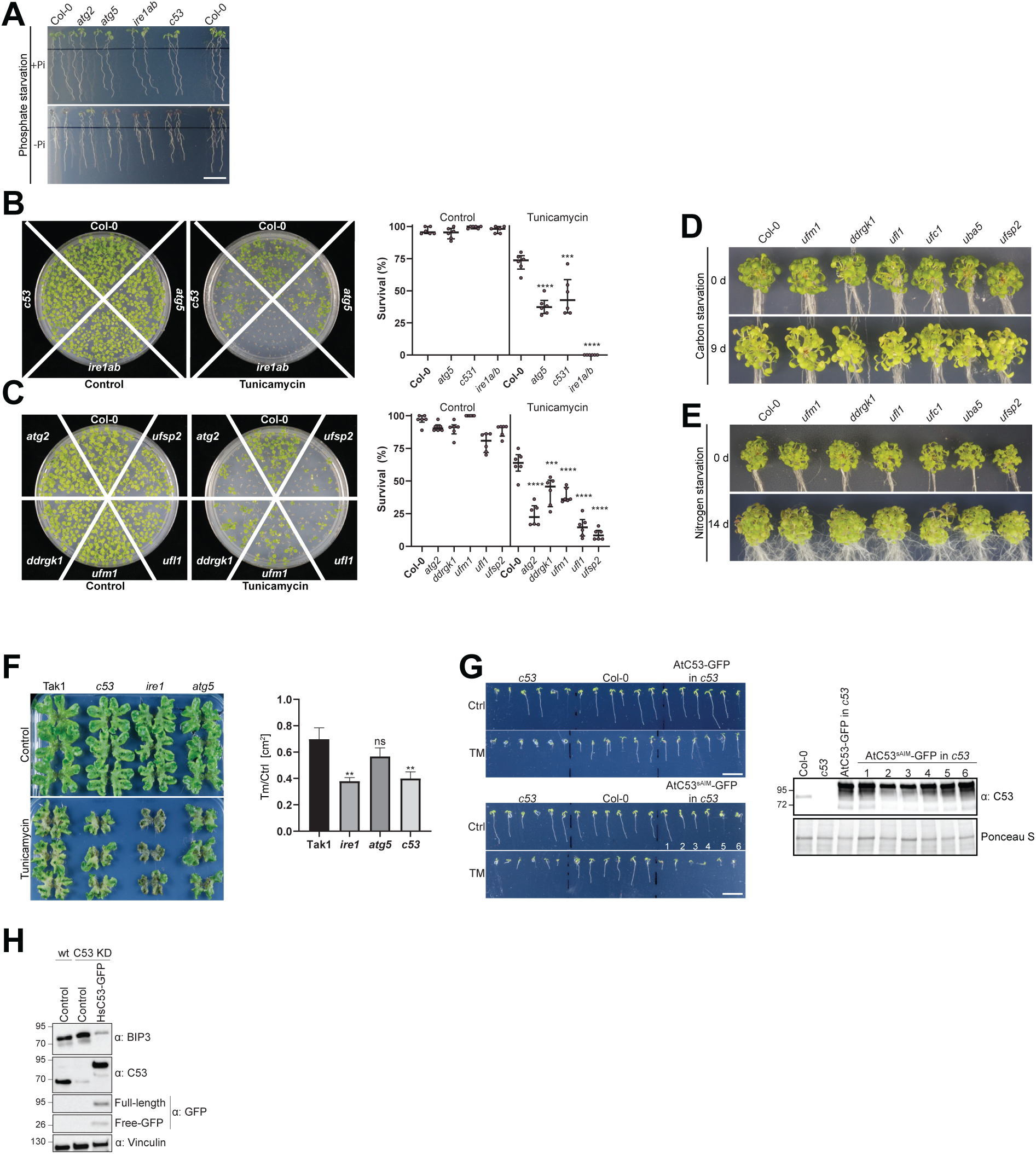
C53 and the UFMylation machinery are essential for ER stress tolerance. **(a) *Atc53* mutant is sensitive to phosphate starvation.** Seedlings were germinated for 5 days on +Pi media prior to transfer for 2 days to medium supplemented with or without Pi (-Pi, +Pi) and imaged after 2 days (Naumann *et al*., 2019). See Figure 5c for quantification. Scale bars, 1 cm. **(b, c) *Atc53* and UFMylation pathway mutants display lower survival on tunicamycin-containing plates.** Seedlings were grown on control or 150 ng/mL tunicamycin containing half-strength MS media for two weeks and were evaluated for survival. *Left Panel* shows an example of the plates. *Right Panel*, Quantification of living and dead seedlings. Data represents 6 plates with 4 x 36 seedlings (n=216). Survival percentage was compared pairwise to the survival of the wildtype (Col-0). Data represent the median with its interquartile range. **(d,e) Ufmylation mutants are insensitive to carbon and nitrogen starvation**. **(d)** Phenotypes before (0 d) and after 9 days carbon starvation (9 d). n = 25. 7-day old seedlings per genotype were used. **(e)** Phenotypes before (0 d) and after 14 days nitrogen starvation (14 d). n = 25. 7-day old seedlings per genotype were used. **(f) *Marchantia polymorpha c53* mutants are sensitive to tunicamycin.** 14-day old plants were transferred to control or 2 µg/ml tunicamycin containing agar plates. *Right Panel,* Quantification of the plant area 21 days after transfer. Data shown as mean ±SD. **(g) The AtC53^sAIM^ mutant does not complement tunicamycin sensitivity phenotype.** *Left Panel,* Representative images of 7-day old seedlings grown on half strength MS media without sucrose in control conditions (Ctrl) or treated with 100 ng/mL tunicamycin (Tm). At least 10 independent T1 transgenic lines were analysed to measure the root lengths. See Figure 5g for quantification. Significant differences compared to control treatment (Ctrl) are indicated with * when p value = 0.05, ** when p value = 0.01, and *** when p value = 0.001. *Right Panel*, western blot analyses of AtC53 complementation lines showing expression of the tested constructs used in Figure 5g. **(h) Knockdown of C53 leads to increased BiP3 chaperone levels.** Knock down and complementation assays showing C53 function is important for ER stress tolerance in HeLa cells.

## Methods

### Cloning procedures

Constructs for *Arabidopsis thaliana* and *£. coli* transformation were generated using the GreenGate (GG) cloning method (Lampropoulos *et al*., 2013). Plasmids used are listed in materials section. The coding sequence of genes of interest were either ordered from Twist Biosciences or Genewiz or amplified from Col-0 or HeLa cDNA using the primers listed in the materials section. The internal BsaI sites were mutated by site-directed-mutagenesis without affecting the amino acid sequence.

For *Marchantia polymorpha* Gateway Cloning (Ishizaki *et al*., 2015) was used to generate all constructs. For HeLa expression experiments, plasmids used are listed in the materials section. The constructs were made by conventional restriction enzyme-based cloning.

### CRISPR/Cas9 construct design

The CRISPR/Cas9 constructs for mutating *c53*, *DDRGKJ* and *UFMJ* in *Arabidopsis thaliana* were prepared according to the protocol described by Xing *et al*., 2014 (Xing *et al*., 2014) and Wang *et al*., 2015 (Ma *et al*., 2015). The pHEE401E and pCBC-DT1T2 vectors for expressing two sgRNAs were provided by Youssef Belkhadir and Jixiang Kong, GMI Vienna. sgRNA target sites were chosen using the website http://crispr.hzau.edu.cn/CRISPR2/. Each gene was targeted by two sgRNAs to remove a fragment of the gene. The CRISPR cassettes of each gene were generated by PCR amplification using pCBC-DT1T2 as template with the primer pairs BsF/F0 and BsR/R0, using adaptors containing the BsaI-restriction sites respectively (see materials section). The PCR products were digested with *Bsa*I, ligated into the pHEE401E plasmid, and transformed into DH5a *£. coli*. Floral dipping proceeded as described previously (Clough and Bent, 1998). Genotyping primers P1 5’-xxx-3’ and P2 5’-xxx-3’ flanking each target site were used to select T1 plants that carried deletions. Sanger sequencing was performed to define the deletion. Through backcrossing with Col-0 plants and genotyping, Cas9-free plants were achieved.

In *Marchantia polymorpha*, CRISPR/Cas9 constructs were generated by selecting two target sequences in *c53* and *ireJ*. Synthetic oligonucleotides were annealed and inserted at the BsaI site of the entry vector pMpGE_En03 flanked by attL1 and attL2 sequences (Sugano *et al*., 2018). The resultant cassettes were inserted to the destination vector pMpGE011 by LR Clonase II Enzyme Mix. The vectors were introduced into thalli of TAK1 via *A. tumefaciens* GV3101+pSoup, and the transformants were selected with 0.5 µM chlorsulfuron (KUBOTA *et al*., 2013). Genomic DNA from transformants was amplified by PCR and sent for sequencing to verify mutations.

### Plant materials and Growth conditions

All *Arabidopsis thaliana* lines used originate from the Columbia (Col-0) ecotype background. Mutant lines used in this study are listed in the materials section. All transgenic plants were generated by the floral dipping method (Clough and Bent, 1998) for which the plasmid constructs were prepared using the green gate cloning method (Lampropoulos *et al*., 2013).

Seeds were then spread on plates or liquid culture with half-strength MS media (Murashige & Skoog salt + Gamborg B5 vitamin mixture) with 1 % sucrose, 0.5 g/L MES and 1% plant agar. pH was adjusted to 5.7 with NaOH. Seeds were imbibed for 4 days at 4 °C in darkness. Plants were grown at 21 °C at 60 % humidity under LEDs with 50 µM/m^2^s and 12h:12h photoperiod.

For the autophagy flux assay, TMT and *in-vivo* immunoprecipitation, seedlings were grown in liquid culture under continuous light.

Male *Marchantia polymorpha* accession Takaragaike-1 (Tak-1) was maintained asexually and cultured through gemma using half-strength Gamborg’s B5 medium containing 1% agar under 50-60 mmol photons m^-2^s^-1^ continuous white light at 22° C unless otherwise defined (KUBOTA *et al*., 2013).

### Plant Sensitivity Tests

#### Arabidopsis thaliana

##### Root-length quantification

Seedlings were grown for 9 days on media supplemented with the indicated drug concentration. Plates were scanned on day 0 and then quantified daily starting from day 2 to day 9. Large scale root-length quantification was conducted using the automated plant imaging analysis software BRAT (Buschlab Root Analysis Toolchain) (Slovak *et al*., 2014) with the inhouse high-performance computer cluster MENDEL. Before analysis, collected data was passed through software quality control.

##### Starvation treatments

Carbon Starvation: Seedlings were grown on half-strength MS media with 1% sucrose for 7 days. They were then transferred to media without sucrose, followed by wrapping the plates in aluminium foil and placing them under the same growth conditions as before for 9 days.

Nitrogen Starvation: Seedlings grew on half-strength MS media with 0.5% sucrose for 7 days. They were then transferred to media without nitrogen and put under the same growth conditions as before for 14 days. Seedlings were arranged in a similar fashion to Jia *et al*., 2019 (Jia *et al*., 2019).

Phosphate Starvation: The method was previously described by Naumann *et al*., 2018 (Naumann *et al*., 2019). Seeds were surface-sterilized and germinated 5 days on +Pi medium prior to transfer to 1% (w/v) Phyto-Agar (Duchefa) containing 2.5 mM KH_2_PO_4_, pH 5.6 (high Pi or +Pi medium) or no Pi supplement (low Pi or -Pi medium), 5 mM KNO_3_, 0.025 mM Fe-EDTA, 2 mM MgSO_4_, 2 mM Ca(NO3)2, 2.5 mM MES-KOH, 0.07 mM H3BO3, 0.014 mM MnCl_2_, 0.01 mM NaCl, 0.5 µM CuSO_4_, 1 µM ZnSO_4_, 0.2 µM Na_2_MoO_4_, 0.01 µM CoCl_2_, 5 g/L sucrose. The agar was routinely purified by repeated washing in deionized water and underwent subsequent dialysis using DOWEX G-55 anion exchanger (Ticconi *et al*., 2009). ICP-MS analysis of the treated agar (7.3 µg/g Fe and 5.9 µg/g P) indicated a contribution of 1.25 µM Fe and 1.875 µM P to the solid 1% agar medium. Images were analysed using ImageJ software.

##### Survival Assay

Seedlings were grown on 9 cm round plates supplemented with the indicated drug at the indicated concentration. Seedling survival was quantified after 14 days. Differentiation between live and dead seedlings was carried out similar to Yang *et al*., 2016 (Yang *et al*., 2016). Surviving seedlings were defined as seedlings which had two green cotyledons and two green true leaves. Plants with yellow leaves or cotyledons were defined as dead.

#### Marchantia polymorpha

##### Tunicamycin Sensitivity

14 days old plants were transformed to half-strength Gamborg’s B5 medium containing indicated concentration of tunicamycin and grown in continues light at 22°C to determine survival rates.

### Autophagy flux assay in *Arabidopsis thaliana*

20-30 seedlings for western blot or 0.5 - 1 g seedlings for immunoprecipitation and mass spectrometry were grown in liquid culture for 5 d under continuous light with shaking at 80 rpm. Media was supplemented with different drugs (3 µM Torin, 10 µg/ml Tunicamycin or other drugs dissolved in DMSO) as indicated. 1 µM of concanamycin was added, if indicated in figures, to track the contribution of vacuolar degradation. For nutrient starvation, seedlings were transferred to phosphate, nitrogen- or sucrose-depleted media (-C, -P, -N). The plants were kept in the dark to reduce sucrose production by photosynthesis or to provide drug stability. Pure DMSO was added to control samples. For analysing total protein degradation such as TMT, seedlings were flash frozen in liquid nitrogen after 24 h treatment. For interaction analysis such as Co-immunoprecipitation, seedling treatment was stopped after 8 h of treatment.

Samples were homogenized in a bead mill (RetschMM300, Haan, Germany; 30 Hz, 90 s) at 4°C with zirconium oxide grinding beads or ground by mortar and pestle for bigger sample volumes. For Western Blotting, SDS loading buffer was added and the sample boiled at 95°C for 10 min. Lysates were cleared by centrifugation at 16,000 g for 10 min and protein concentration was normalized by Amidoblack staining (Sigma). Western blotting was performed following standard protocols as described below. 5 µg of lysate was loaded per lane.

### Human Cell Culture conditions

HeLa-Kyoto and HEK293T cells maintained in Dulbecco’s modified Eagle’s Medium (DMEM) with 10% FBS, 1% L-Glutamine and 1% Penicillin/Streptomycin. Transfection was performed with GeneJuice transfection reagent according to manufacturer’s instructions. 100 µl of empty media was mixed with 3µL of GeneJuice and after 5 min of incubation a total of 1 µg of DNA mixture per transfection was added. After 20 minutes of incubation transfection mixture was added dropwise to the cells. Cells were incubated with DNA for 24h. DNA containing media was removed and replaced with media.

### Lentiviral Knockdown

#### Lentiviral transduced shRNA mediated knockdown of *c53* in HeLa cells

The knockdown was performed in S2 conditions. HEK293T cells were seeded 24 h prior to transfection in DMEM without antibiotics. At 50-60 % confluency, cells were transfected with 1 µg shRNA, 750 ng psPAX2 and 250 ng pMD2.G utilizing 6 µL of GeneJuice in 250 µL of empty DMEM. After 48 h of incubation the virus containing media was harvested and mixed 1:1 with full media. This mixture was applied to HeLa cells that were seeded 24 h prior. Polybrene was added to a final concentration of 4 µg/ml. After 24 h of incubation the medium on target cells was exchanged with full media. After 24h, selection with 2 µg/ml Puromycin was started. No living cells were observed in a control plate after 24 h. After splitting cells in S2 conditions, cells were transferred into S1 conditions.

### Autophagy Flux Assay in Human cell culture

Cells were seeded 24 h prior to treatment. At 50-60 % confluency treatments were started by replacing media containing the indicated drugs or full media (untreated). Tunicamycin was added with a final concentration of 2.5 µg/ml and Torin with a final concentration of 3 µM. The treatments were stopped after 16h by removing the media and washing the cells with 1xPBS. A 2 h recovery period was started by adding either media containing 100 nM Bafilomycin A1 or full media. Cells were put on ice and lysed with 100 µL of Lysis buffer (50 mM HEPES, 150 mM NaCl, 1 mM EDTA, 1 mM EGTA, 25 mM NaF, 10 µM ZnCl_2_, 1% Triton X-100 and 10% Glycerol) per well. After centrifugation, supernatant was mixed 1:1 with 2x Laemmli Buffer and denatured by heating to 95°C for 5min.

Each sample was loaded onto a 4-20% SDS-PAGE gradient gel (BioRad) and electrophoresis was run at 100V for 1,5h.

### Western Blotting

SDS-PAGE was performed using gradient 4-20% Mini-PROTEAN® TGX Precast Protein Gels (BioRad). Blotting on nitrocellulose membranes was performed using a semi-dry Turbo transfer blot system (BioRad). For images of human LC3B, a wet transfer to PVDF membranes was performed at 200mA for 70 minutes. Membranes were blocked with 5% skimmed milk or BSA in TBS and 0.1% Tween 20 (TBS-T) for 1h at room temperature or at 4°C overnight. This was followed by incubation with primary and subsequent secondary antibody conjugated to horseradish peroxidase. After 3 times 10 min washes with TBS-T, the immune-reaction was developed using ECL Super-Pico Plus (Thermo) and detected with ChemiDoc Touch Imaging System (BioRad).

### Western Blot Image Quantification

Protein bands intensities were quantified with Image Lab 6 (BioRad). Equal rectangles were drawn around the total protein gel lane and the band of interest. The lane profile was obtained by subtracting the mean intensity of the background. The adjusted volume of the peak in the profile was taken as a measure of the band intensity. The protein band of interest was normalized for the total protein level of the whole lane. Average relative intensities and a standard error of at least 3 independent experiments were calculated.

### Chemicals and Antibodies

To generate *At*C53 antibody, purified protein was sent to Eurogentec for immunization of rabbits via their 28-day program. The final bleed was purified on column conjugated with the purified protein.

### *In-vitro* Pulldowns

For pulldown experiments, 10 µl of glutathione magnetic agarose beads (Pierce™ Glutathione Magnetic Agarose Beads, Thermo Scientific™) were equilibrated by washing them 2 times with wash buffer (100 mM Sodium Phosphate pH 7.2, 300 mM NaCl, 1 mM DTT, 0.01% (v/v) IGEPAL). Normalized *£. coli* clarified lysates or purified proteins were mixed, according to the experiment, added to the washed beads and incubated on an end-over-end rotator for 1hr at 4°C. Beads were washed 5 times in 1 ml wash buffer. Bound proteins were eluted by adding 100 µl Laemmli buffer. Samples were analysed by western blotting or Coomassie staining.

### Yeast two hybrid assay (Y2H)

Yeast two hybrid assay (Y2H) was performed according to the Mathmaker^TM^ GAL4 Two hybrid system (Clonetech®) following the protocol from the manufacture. Different genes were fused in frame to GAL4 activation domain of the prey vector pGADT7 and GAL4 binding domain from the bait vector pGBKT7. Split-GFP was used as positive control. Combinations of pGADT7 and pGBKT7 vectors carrying the different genes were transformed in the yeast strains Y187 (MAT a) and AH109 (MAT a), respectively. After mating between bait and prey strains, the diploid yeast was selected for growth on (SD)-Leu/-Trp, (SD)-Leu/-Trp/-His and (SD)-Leu/-Trp/-His/-Ade plates at 28°C for 2 to 4 days.

### *In planta* Co-Immunoprecipitation

0.5 - 1 g seedlings were grown in liquid and treated as described under section Autophagy Flux Assay. After grinding of frozen samples, G-TEN buffer (10% Glycerole, 50 mM Tris/HCl pH 7.5, 1mM EDTA, 300 mM NaCl, 1 mM DTT, 0.1% [v/v] Nonidet P-40/Igepal, Complete protease inhibitor tablet) was added, vortexed, and lysates were cleared by centrifugation at 16 000 g for 10 min at 4°C. Protein concentration was equally adjusted using Bradford protein assay (Sigma).

25 µl of GFP-Trap_A beads (Chromotek) were equilibrated and added to each lysate and incubated for 2 h at 4°C on a turning wheel. Beads were washed 3 times with 1 mL G-TEN buffer.

For Western Blot analysis, beads were resuspended in 30 µl SDS-loading buffer (116 mM Tris-HCl pH 6.8, 4.9% glycerol, 10 mM DTT, 8% SDS). On-bead bound proteins were eluted by boiling the beads for 10 min at 70 °C and analysed by western blotting with indicated antibodies.

For mass spectrometry experiments, the beads were further washed 5 times with mass spectrometry compatible buffer (50 mM Tris/HCl pH 7.5, 1mM EDTA). Buffer resuspended beads were then submitted for trypsin digestion.

### Microscopy-based protein-protein interaction assays

Bead-bound bait proteins were incubated with fluorescently labelled prey protein as described previously by Turco *et al.* 2019 (Turco *et al*., 2019). 10 µl of Glutathione Sepharose 4B beads (GE Healthcare, average diameter 90mm) were incubated for 30 min at 4 °C (16 rpm horizontal rotation) with GST-tagged bait proteins (4 mg/mL for GST and GST-FIP200 CTR). The beads were washed 2 times in 10x bead volume of washing buffer (25 mM HEPES pH 7.5, 150 mM NaCl, 1 mM DTT). The buffer was removed, and the beads were resuspended 1:1 in washing buffer. 10 µL of a 2-5 µM dilution of fluorescently labelled binding partners (GFP, C53-GFP and GFP-p62) were added to the bead suspension and incubated for 30-60 min at room temperature before imaging with a Zeiss LSM700 confocal microscope with 20 X magnification. For Quantification the maximum gray value along the diameter of each bead (n = 15) was measured.

### Mass Spectrometry (TMT) and Analysis

MS/MS Data Analysis: Raw files were processed with Proteome Discoverer (version 2.3, Thermo Fisher Scientific, Bremen, Germany). Database searches were performed using MS Amanda (version 2.3.0.14114) (Dorfer *et al*., 2014) against the TAIR10 database (32785 sequences). The raw files were loaded as fractions into the processing workflow. Carbamidomethylation of cysteine and TMT on peptide N-termini were specified as fixed modifications, phosphorylation on serine, threonine and tyrosine, oxidation of methionine, deamidation of asparagine and glutamine, TMT on lysine, carbamylation on peptide N-termini and acetylation on protein N-termini were set as dynamic modifications. Trypsin was defined as the proteolytic enzyme, cleaving after lysine or arginine. Up to two missed cleavages were allowed. Precursor and fragment ion tolerance were set to 5 ppm and 15 ppm respectively. Identified spectra were rescored using Percolator (Kall *et al*., 2007), and filtered to 0.5% FDR at the peptide spectrum match level. Protein grouping was performed in Proteome Discoverer applying strict parsimony principle. Proteins were subsequently filtered to a false discovery rate of 1% at protein level. Phosphorylation sites were localized using IMP-ptmRS implemented in Proteome Discoverer using a probability cut-off of >75% for unambiguous site localization.

TMT-quantification: TMT Reporter ion S/N values were extracted from the most confident centroid mass within an integration tolerance of 20 ppm. PSMs with average TMT reporter S/N values below 10 as well as PSMs showing more than 50% co-isolation were removed. Protein quantification was determined based on unique peptides only. Samples were sum normalized and missing values were imputed by the 5% quantile of the reporter intensity in the respective sample. Statistical significance of differentially abundant proteins was determined using limma (Smyth, 2004). Gene Ontology (Ashburner *et al*., 2000) enrichment was determined using DAVID (Dennis *et al*., 2003) (version 6.8). Cross species comparison of regulated proteins was performed by mapping proteins to ortholog clusters available in eggnog (Huerta-Cepas *et al*., 2015). Proteins containing signal peptides were predicted using SignalP 5.0 (Almagro Armenteros *et al*., 2019).

### Peptide array

High density peptide array analysis was performed commercially by PEPperPRINT. This comprised a full substitution scan of wild type peptide GVSEWDPILEELQEM, with exchange of all amino acid positions with 23 amino acids including citrulline (Z), methyl-alanine (O) and D-alanine (U). The analysis also included an N- and C-terminal deletion series of wild type peptide GVSEWDPILEELQEM; an additional 32 spots of custom control peptide KPLDFDWEIVLEEQ, and acidic variants of this control peptide involving exchanges of selected amino acid positions with glutamic acid €. The resulting peptide microarrays contained 416 different linear peptides printed at least in triplicate (1,412 peptide spots; wild type peptides were printed with a higher frequency), and were framed by HA (YPYDVPDYAG, 88 spots) control peptides (See Table S1 for the array map).

Peptide microarrays were pre-stained with rabbit anti-GST Dylight680 at a dilution of 1:2000 to investigate background interactions with the variants of wild type peptides GVSEWDPILEELQEM and KPLDFDWEIVLEEQ that could interfere with the main assays. Subsequent incubation of other peptide microarrays with proteins GST-ATG8A and GST at a concentration of 10 µg/ml in incubation buffer was followed by staining with secondary antibody rabbit anti-GST Dylight680 and read-out at a scanning intensity of 7 (red). The control staining of the HA epitopes with control antibody mouse monoclonal anti-HA (12CA5) DyLight800 was finally done as an internal quality control to confirm the assay quality and the peptide microarray integrity. Read-out of the control staining was performed at a scanning intensity of 7/7 (red/green).

Quantification of spot intensities and peptide annotation were based on the 16-bit grey scale tiff files at a scanning intensity of 7 that exhibit a higher dynamic range than the 24-bit colorized tiff files; microarray image analysis was done with PepSlide® Analyzer. A software algorithm breaks down fluorescence intensities of each spot into raw, foreground and background signal (see “Raw Data” tabs), and calculates averaged median foreground intensities and spot-to-spot deviations of spot duplicates.

### Microscopy Methods

#### Preparation of *Arabidopsis thaliana* samples for confocal imaging

4d old seedlings were treated as indicated under the autophagy flux assay section. Seedlings were imaged between 3h - 6h of drug incubation. Roots were placed on a microscope slide with indicated treatment buffer and closed with coverslip. Imaging was performed in the root differentiation zone where root hair growth starts.

##### Preparation of human cell samples for confocal imaging

Transfected and treated cells were grown on coverslips and fixed utilizing 0.4% Paraformaldehyde solution in PBS for 30min. Fixed cells were mounted in VectaShield mounting medium without DAPI and sealed using clear nail polish.

#### Confocal imaging

Samples were imaged at an upright ZEISS LSM800 or LSM 780 confocal microscope (Zeiss) with an Apochromat 40x or 63x objective lens at 1x magnification.

Excitation/detection parameters for GFP and mCherry were 488 nm/463 nm and 510 nm and 561 nm/569 to 635 nm respectively, and sequential scanning mode was used for colocalization of both fluorophores. Identical settings, including an optical section thickness of 2 µm per z-stack, were used during the acquisition for sample comparison, and the images processed using identical parameters. Confocal images were processed with ZEN (version 2011) and ImageJ (version 1.48v) software.

#### Image Quantification

Autophagic puncta were counted using ImageJ. Several (at least five) z-stack merged images were manually background subtracted, thresholded and the same threshold value was applied to all the images and replicates of the same experiment. The image was converted to eight-bit grayscale and then counted for ATG8 puncta either manually or by the Particle Analyzer function of ImageJ. The average number of autophagosomes per z-stack was averaged between 10 or more different roots.

Colocalization analysis was performed by calculating Pearson’s correlation coefficient as previously described using ImageJ software with the plug-in JACoP (BOLTE and CORDELIERES, 2006). Values near 1 represent almost perfect correlation, whereas values near 0 reflect no correlation. The average Pearson’s correlation coefficient was determined in 5 or more different roots.

#### Ultrastructural analyses using immunogold labelling electron microscopy

##### TEM experiments using mCherry and native AtC53 antibodies

For high pressure freezing, 5-day old *Arabidopsis* seedling roots expressing AtC53-mCherry were cut and high-pressure frozen (EM PACT2, Leica, Germany), prior to subsequent freeze substitution in acetone containing 0.4% uranyl acetate at −85 °C in an AFS freeze-substitution unit (Leica, Wetzlar, Germany). After gradient infiltration with increasing concentration of HM20, root samples were embedded and ultraviolet polymerized for ultra-thin sectioning and imaging. TEM images were captured by an 80 kV Hitachi H-7650 transmission electron microscope (Hitachi High-Technologies Corporation, Japan) with a charge-coupled devise camera. IEM analysis were performed as previously described (Zhuang *et al*., 2017).

##### TEM experiments using GFP antibodies

*Arabidopsis* roots were fixed in 2% paraformaldehyde and 0.2% glutaraldehyde (both EM-grade, EMS, USA) in 0.1 M PHEM buffer (pH 7) for 2h at RT, then overnight at 4°C. The fixed roots were embedded in 12% gelatin and cut into 1 mm^3^ blocks which were immersed in 2.3 M sucrose overnight at 4°C. These blocks were mounted onto a Leica specimen carrier (Leica Microsystems, Austria) and frozen in liquid nitrogen. With a Leica UCT/FCS cryo-ultramicrotome (Leica Microsystems, Austria) the frozen blocks were cut into ultra-thin sections at a nominal thickness of 60nm at −120°C. A mixture of 2% methylcellulose (25 centipoises) and 2.3 M sucrose in a ratio of 1:1 was used as a pick-up solution. Sections were picked up onto 200 mesh Ni grids (Gilder Grids, UK) with a carbon coated formvar film (Agar Scientific, UK). Fixation, embedding and cryo-sectioning was conducted as described by Tokuyasu *et al*., 1973.

###### Immunolabeling

Prior to immunolabeling, grids were placed on plates with solidified 2% gelatine and warmed up to 37 °C for 20 min to remove the pick-up solution. After quenching of free aldehyde-groups with glycine (0.1% for 15 min), a blocking step with 1% BSA (fraction V) in 0.1 M Sorensen phosphate buffer (pH 7.4) was performed for 40 min. The grids were incubated in primary antibody, rabbit polyclonal to GFP (ab6556, Abcam, UK), diluted 1:125 in 0.1 M Sorensen phosphate buffer over night at 4°C, followed by a 2h incubation in the secondary antibody, a goat-anti-rabbit antibody coupled with 6 nm gold (GAR 6 nm, Aurion, The Netherlands), diluted 1:20 in 0.1 M Sorensen phosphate buffer, performed at RT. The sections were stained with 4% uranyl acetate (Merck, Germany) and 2% methylcellulose at a ratio of 1:9 (on ice). All labeling steps were conducted in a wet chamber. The sections were inspected using a FEI Morgagni 268D TEM (FEI, The Netherlands) operated at 80kV. Electron micrographs were acquired using an 11-megapixel Morada CCD camera from Olympus-SIS (Germany).

### Statistical analysis

Statistical analyses were performed with GraphPad Prism 8 software. For all the quantifications described above, statistical analysis was performed. Statistical significance of differences between two experimental groups was assessed wherever applicable by either a two-tailed Student’s t-test if the variances were not significantly different according to the F test, or using a non-parametric test (Mann-Whitney or Kruskal-Wallis with Dunn’s post-hoc test for multiple comparisons) if the variances were significantly different (p < 0.05). Differences between two data sets were considered significant at p < 0.05 (*) (p < 0.01 (**); p < 0.001 (***); p < 0.0001 (****); n.s., not significant.

### Biophysical Characterization

#### Protein purification

Recombinant proteins were produced using *£. coli* strain Rosetta2(DE3)pLysS grown in 2x TY media at 37°C to an A600 of 0.4-0.6 followed by induction with 300 µM IPTG and overnight incubation at 18°C. Pelleted cells were resuspended in lysis buffer (100 mM Sodium Phosphate pH 7.0, 300 mM NaCl) containing protease inhibitors (Complete™, Roche) and sonicated. The clarified lysate was first purified by affinity, by using HisTrap FF (GE HealthCare) columns. The proteins were eluted with lysis buffer containing 500 mM Imidazole. The eluted fraction was buffer exchanged to 10 mM Sodium Phosphate pH 7.0, 100 mM NaCl and loaded either on Cation Exchange, Resource S, or Anion Exchange, Resource Q, chromatography columns. The proteins were eluted by NaCl gradient (50% in 20 CV). Finally, the proteins were separated by Size Exclusion Chromatography with HiLoad® 16/600 Superdex® 200 pg or HiLoad® 16/600 Superdex® 75 pg, which were previously equilibrated in 50 mM Sodium Phosphate pH 7.0, 100 mM NaCl. The proteins were concentrated using Vivaspin concentrators (3000, 5000, 10000 or 30000 MWCO). Protein concentration was calculated from the UV absorption at 280 nm by DS-11 FX+ Spectrophotometer (DeNovix).

#### Surface plasmon resonance analysis

Binding of AIM *wt* (EPLDFDWEIVLEEEM) and AIM mutant (EPLDFDAEIALEEEM) peptide to GST-GABARAP and GST-ATG8A, respectively, was investigated by surface plasmon resonance analysis using a Biacore T200 instrument (GE Healthcare) operated at 25°C. In additione, AIM-dependent binding of *Hs*C53 and *At*C53 to GST-GABARAP and GST-ATG8A were studied. The running buffer used for all experiments was 50 mM sodium phosphate pH 7.0 supplemented with 100 mM NaCl, 0.05% (v/v) Tween-20 and 0.1% (w/v) bovine serum albumin.

Polyclonal anti-GST antibodies (GST Capture Kit, GE Healthcare) were amine coupled on to a Series S CM5 sensor chip (GE Healthcare) using 2 adjacent flow cells (i.e. the reference and active cell) according to the manufacturer’s instructions.

To determine specific binding, GST-GABARAP or GST-ATG8A were captured on the active cell (concentration: 5 µg/ml; contact time: 30 seconds; flow rate: 10 µl/min) and GST was captured on the reference cell (concentration: 10 µg/ml; contact time: 30 seconds; flow rate: 10 µl/min) to perform background subtraction.

To qualitatively show whether the analytes, *Hs*C53, *Hs*C53 123A (i.e. HsC53^W269A, W294A, W312A^), *At*C53 and *At*C53 1234A (i.e. *At*C53^W276A, W287A, Y304A, W335A^), interact or do not interact in an AIM-dependent manner with GST-GABARAP or GST-ATG8A, the 2 flow cells were exposed to 4 sets of double consecutive injections (1^st^ set: 10 µM analyte, running buffer; 2^nd^ set: 10 µM analyte, 10 µM analyte; 3^rd^ set: 10 µM analyte, 10 µM analyte + 6.4 µM AIM *wt* peptide; 4^th^ set: 10 µM analyte, 10 µM analyte + 6.4 µM AIM mutant peptide. Contact time 1^st^ injection: 30 seconds; contact time 2^nd^ injection: 30 seconds; dissociation time: 60 seconds; flow rate: 30 µl/min).

To quantify the binding affinities of the AIM *wt* peptide to GST-ATG8 or GST-GABARAP, multi-cycle kinetic experiments with increasing concentrations of the AIM *wt* peptide (25, 50, 100, 200, 400, 800, 1600, 3200 nM and 400 nM as internal replicates) were performed (contact time: 60 seconds; dissociation time: 60 seconds; flow rate: 30 µl/min). As a negative control, the chip was exposed to 3200 nM of the AIM mutant peptide (contact time: 60 seconds; dissociation time: 60 seconds; flow rate: 30 µl/min).

To quantify the apparent binding affinity of the AIM *wt* peptide to GST-GABARAP in presence of *Hs*C53, multi-cycle kinetic experiments with increasing concentrations of the AIM peptide (0, 25, 50, 100, 200, 400, 800, 1600, 3200 nM and 400 nM as internal replicates), containing 10 µM of *Hs*C53, in running buffer (contact time: 60 seconds; dissociation time: 60 seconds; flow rate: 30 µl/min). For negative controls, the chip was exposed to 3200 nM of the AIM mutant peptide, containing 10 µM of *Hs*C53 or 10 µM of *Hs*C53 123A, and to 3200 nM of the AIM *wt* peptide, containing 10 µM of *Hs*C53 123A (contact time: 60 seconds; dissociation time: 60 seconds; flow rate: 30 µl/min).

To quantify the apparent binding affinity of the AIM *wt* peptide to GST-ATG8A in the presence of *At*C53, multi-cycle kinetic experiments with increasing concentrations of the AIM peptide (0, 50, 100, 200, 400, 800, 1600, 3200, 6400, 12800 nM and 400 nM as internal replicate) containing 10 µM of *At*C53 were performed (contact time: 60 seconds; dissociation time: 60 seconds; flow rate: 30 µl/min). As negative controls, the chip was exposed to 6400 nM of the AIM mutant peptide, containing 10 µM of *At*C53 or 10 µM of *At*C53 1234A, and to 6400 nM of the AIM *wt* peptide, containing 10 µM of *At*C53 1234A (contact time: 60 seconds; dissociation time: 60 seconds; flow rate: 30 µl/min).

After each cycle, regeneration was performed with 2 injections of 10 mM glycine-HCl pH 2.1 for 120 seconds at a flow rate of 10 µL/min.

The sensograms obtained were analysed with Biacore T200 Evaluation software (version 3.1) by global fitting of the data to a 1:1 steady-state affinity model.

Molecular weights and sources of the proteins for the SPR experiments are reported in the following table:

| Sample name | Source | MW (Da) |
| --- | --- | --- |
| AIM *wt* peptide | Synthesized *in house* | 1894.08 |
| AIM mutant peptide |  | 1750.89 |
| *HsC53* | *Escherichia coli*<br>recombinant<br>expression | 59191.15 |
| *HsC53* 123A |  | 58758.66 |
| *AtC53* |  | 64399.57 |
| *AtC53* 1234A |  | 63962.07 |
| GST-GABARAP |  | 42458.87 |
| GST-ATG8A |  | 42366.85 |
| GST |  | 27898.33 |

Calculation for the apparent *K_D_* (*K′_D_*) of the AIM^wt^ was done by using the following formula (Nelson, David L. Lehninger Principles Of Biochemistry. New York: W.H. Freeman, 2008):

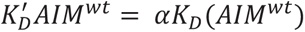

Where, 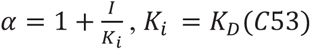 and *[I] = [C53]*

Then:

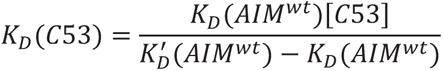

### Isothermal titration calorimetry (ITC)

All experiments were carried out at 25°C in 50 mM sodium phosphate buffer pH 7.0, 100 mM NaCl, using the PEAQ-ITC Automated (Malvern Panalytical Ltd). For protein-protein interactions, the calorimetric cell was filled with 40 µM GABARAP or ATG8A and titrated with 250 µM *Hs*C53 or *At*C53 IDRs, respectively. A single injection of 0.4 µl of *Hs*C53 or *At*C53 IDRs (not taken into account) was followed by 18 injections of 2 µl each. Injections were made at 150 sec intervals with a duration of 4 sec and a stirring speed of 750 rpm. The reference power was set to 10 µcal/sec, the feedback mode was set to *high*. For protein-peptide interactions, the calorimetric cell was filled with 40 µM GABARAP or ATG8A and titrated with 600 µM peptide from the syringe. The titrations were performed as described above. For the control experiments, equivalent volumes of the IDRs, or the peptides, were titrated to buffer, equivalent volumes of buffer were titrated to GABARAP or ATG8A and equivalent volumes of buffer were titrated to buffer, using the parameters above. The raw titration data were integrated, corrected for the controls and fitted to a one-set-of-sites binding model using the PEAQ-ITC analysis software (Version 1.22).

### Sample preparation for native MS experiments

Proteins were buffer exchanged into ammonium acetate using BioRad Micro Bio-Spin 6 Columns and concentrations were measured using DS-11 FX+ Spectrophotometer (DeNovix).

### Mass spectrometry measurements

Native mass spectrometry experiments were carried out on a Synapt G2S*i* instrument (Waters, Manchester, UK) with a nanoelectrospray ionisation source. Mass calibration was performed by a separate infusion of NaI cluster ions. Solutions were ionised through a positive potential applied to metal-coated borosilicate capillaries (Thermo Scientific). The following instrument parameters were used; capillary voltage 1.3 kV, sample cone voltage 40 V, extractor source offset 30 V, source temperature 40 °C, trap gas 3 mL/min. A higher capillary voltage (1.9 kV) was required for ionization of the 1:2 AtC53-AtG8A complex. Data were processed using Masslynx V4.1 and spectra were plotted using R. Peaks were matched to protein complexes by comparing measured m/z values with expected m/z values calculated from the mass of individual proteins which are given in table below.

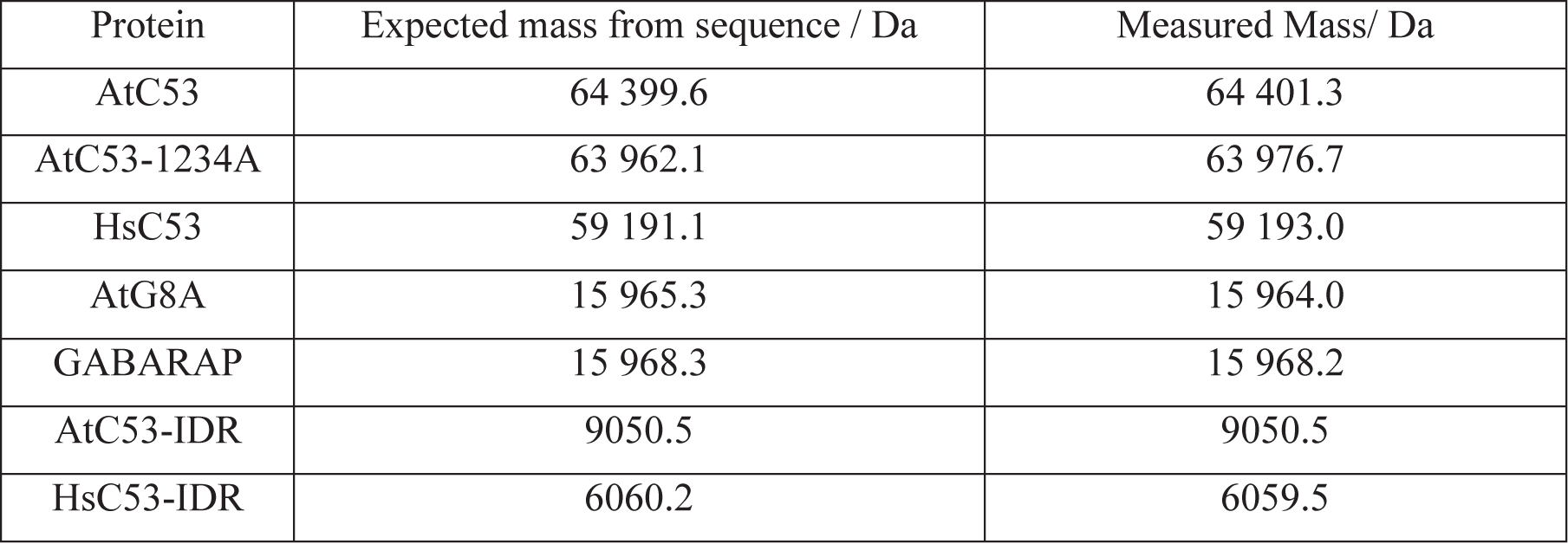

### Circular dichroism spectroscopy

CD spectroscopy experiments were performed using a Chirascan-Plus CD spectrophotometer (Applied Photophysics). Purified proteins in 50mM sodium phosphate pH 7.0, 100mM NaCl were diluted to approximately 0.2 mg/ml and spin-filtered with an 0.1µm filter. CD measurements were carried out in a quartz glass cuvette with 0.5 mm path length. To obtain overall CD spectra, wavelength scans between 180nm and 260nm were collected at 25°C using a 1.0 nm bandwidth, 0.5 nm step size, and time per point of 0.5 s. Both CD and absorbance data were collected at the same time over three accumulations and averaged. CD data at wavelengths where the absorptivity was above 2.5 are not shown (data below 194nm). The raw data in millidegree units were corrected for background and drift (*e_dcorr_*). Subsequently the differential molar extinction coefficient per peptide bond (11s) was calculated, taking into account the absorptivity measured at 205nm (A_205_) and the calculated protein extinction coefficient at 205nm (s_205_) using the equation

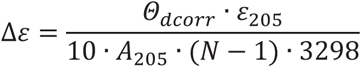

## Materials

### Experimental Model/Cell lines

| Organism | Background | Supplier |
| --- | --- | --- |
| *Arabidopsis thaliana* | Col-0 |  |
| *Marchantia Polymorpha* | Tak-1 |  |
| Human Tumor Cell line | HeLa-Kyoto | Fumiyo Ikeda |
| Human Embryonic Kidney Cells | HEK293T | Fumiyo Ikeda |

### Plant Mutants

| Genotype | Gene ID | Preparation/T-DNA line | Supplier | Reference |
| --- | --- | --- | --- | --- |
| *c53* | At5g06830 | CRISPR/Cas9 | this study |  |
| *ddrgk1* | At4g27120 | CRISPR/Cas9 | this study |  |
| *ufm1* | At1g77710 | CRISPR/Cas9 | this study |  |
| *atg2* | At3g19190 | EMS-mutant (Gln803stop) | Morten Peterson | Wang *et al.* Plant Journal (2011) |
| *atg5* | At5g17290 | SAIL_129B07 | NASC (N39993) | Scholl *et al.* Plant Phys. (2000) |
| *ufl1* | At3g46220 | SALK_022517C | NASC (N685434) | Scholl *et al.* Plant Phys. (2000) |
| *uba5* | At1g05350 | SALK_134012 | NASC (N634012) | Scholl *et al.* Plant Phys. (2000) |
| *ufsp2* | At3g48380 | SAIL_607_G10 | NASC (N826004) | Scholl *et al.* Plant Phys. (2000) |
| *ufc1* | At1g27530 | SALK_112532 | NASC (N678973) | Scholl *et al.* Plant Phys. (2000) |
| *ire1a/b* | At2G17520/<br>At5G24360 | SALK_018112/SAIL_238_F07 | Karolina Pajeroska-Mukhtar | McCormack *et al.* Front. in plant sci. (2015) |
| *bzip 17/28* | At2g40950/<br>At3g10800 | SALK_104326/ SALK_132285 | Kazuo Shinozaki | Kim *et al.* Plant Phys. (2018) |

### Plant Lines (all Col-0 background)

| Name | Selection | Supplier | Reference |
| --- | --- | --- | --- |
| pUbi::mCherry-ATG8A | Alli-mCherry | this study |  |
| pUbi::mCherry-ATG8E | HygB | Liwen Jiang | Hu *et al.* J. Integr. Plant Biol. (2020) |
| pUbi::mCherry-ATG8E x *atg5* | HygB/Basta | this study |  |
| pUbi::GFP-ATG8A | Alli-YFP | this study |  |
| pUbi::GFP-ATG8B | Alli-YFP | this study |  |
| pUbi::GFP-ATG8C | Alli-YFP | this study |  |
| pUbi::GFP-ATG8D | Alli-YFP | this study |  |
| pUbi::GFP-ATG8E | Alli-YFP | this study |  |
| pUbi::GFP-ATG8F | Alli-YFP | this study |  |
| pUbi::GFP-ATG8G | Alli-YFP | this study |  |
| pUbi::GFP-ATG8H | Alli-YFP | this study |  |
| pUbi::GFP-ATG8I | Alli-YFP | this study |  |
| pUbi::C53-mCherry | Kan | this study |  |
| pUbi::C53-mCherry x *atg2* | Kan | this study |  |
| pUbi::C53-mCherry x *atg5* | Kan | this study |  |
| pUbi::C53-mCherry x *ufl1* | Kan | this study |  |
| pUbi::C53-mCherry x *ddrgk1* | Kan | this study |  |
| pUbi::C53-mCherry x *ire1a/b* | Kan | this study |  |
| pUbi::C53-mCherry x pUbi::GFP-ATG8A | Alli-YFP/Kan | this study |  |
| pUbi::C53-mCherry x wave-YFP | Kan/Bas | this study |  |
| pUbi::C53-mCherry x p35S::GFP-HDEL | Kan | this study |  |
| pUbi::C53-mCherry x p35S::GFP-ATG11 | Kan/HygB | this study |  |
| pUbi::C53-mCherry x pUbi::UFL1-GFP | Kan/Alli-YFP | this study |  |
| pUbi::C53-mCherry x pUbi::DDR GK1-GFP | Kan/Alli-YFP | this study |  |
| pUbi::C53-mCherry x GCSI-SUBEX-C57Y-GFP | Kan/HygB | this study/<br>Richard Strasser | Shin *et al.* The Plant Journal (2018) |
| pUbi::C53-mCherry x MNS1-SUBEX-GFP | Kan/HygB | this study/<br>Richard Strasser | Shin *et al.* The Plant Journal (2018) |
| pUbi::C53-GFP | Alli-YFP | this study |  |
| pUbi::C53-GFP x *c53* | Alli-YFP | this study |  |
| pUbi::C53<sup>sAIM(W276A,W287A,Y304A,W335A)</sup>-GFP x *c53* | Alli-YFP | this study |  |
| pUbi::C53-GFP x SP-mRFP-SUBEX-C57Y-EMP12 | Alli-YFP/HygB | this study/<br>Richard Strasser | Shin *et al.* Plant Journal (2018) |
| pUbi::UFL1-GFP | Alli-YFP | this study |  |
| pUbi::DDR GK1-GFP | Alli-YFP | this study |  |
| p35S::GFP-HDEL (ER-gk) | Kan | NASC | N16251 |
| wave-YFP | BASTA | Niko Geldner | Geldner et al. The Plant Journal (2009) |

### Bacterial Strains

| Organism | Strain | Supplier |
| --- | --- | --- |
| *E. coli* | DH5α | In-house facility |
|  | BL21 (DE3) | In-house facility |
|  | Rosetta2 (DE3) pLysS | In-house facility |
| *Agrobacterium tumefaciens* | GV3101 (pSoup) | In-house facility |

### Oligonucleotides - gRNAs

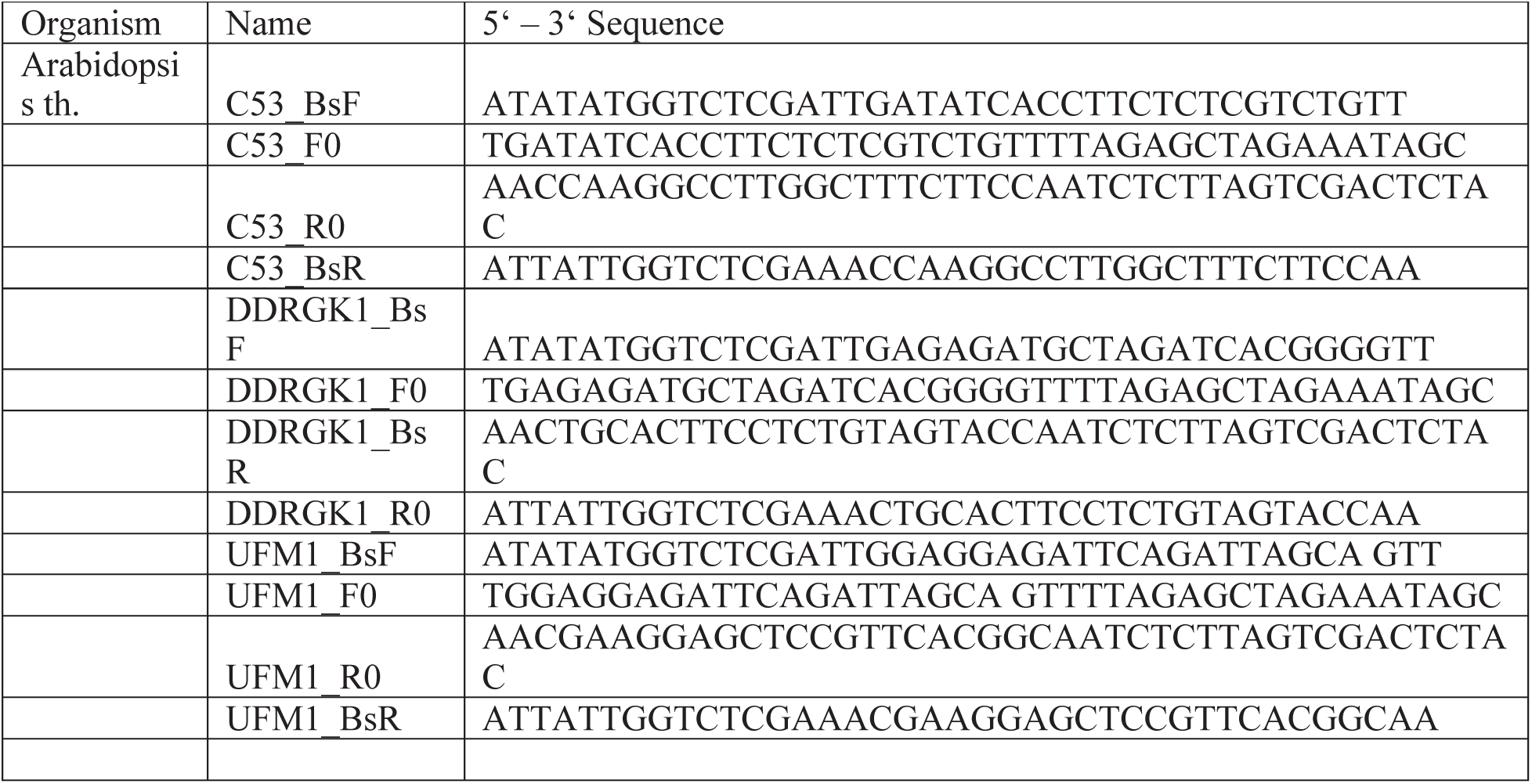
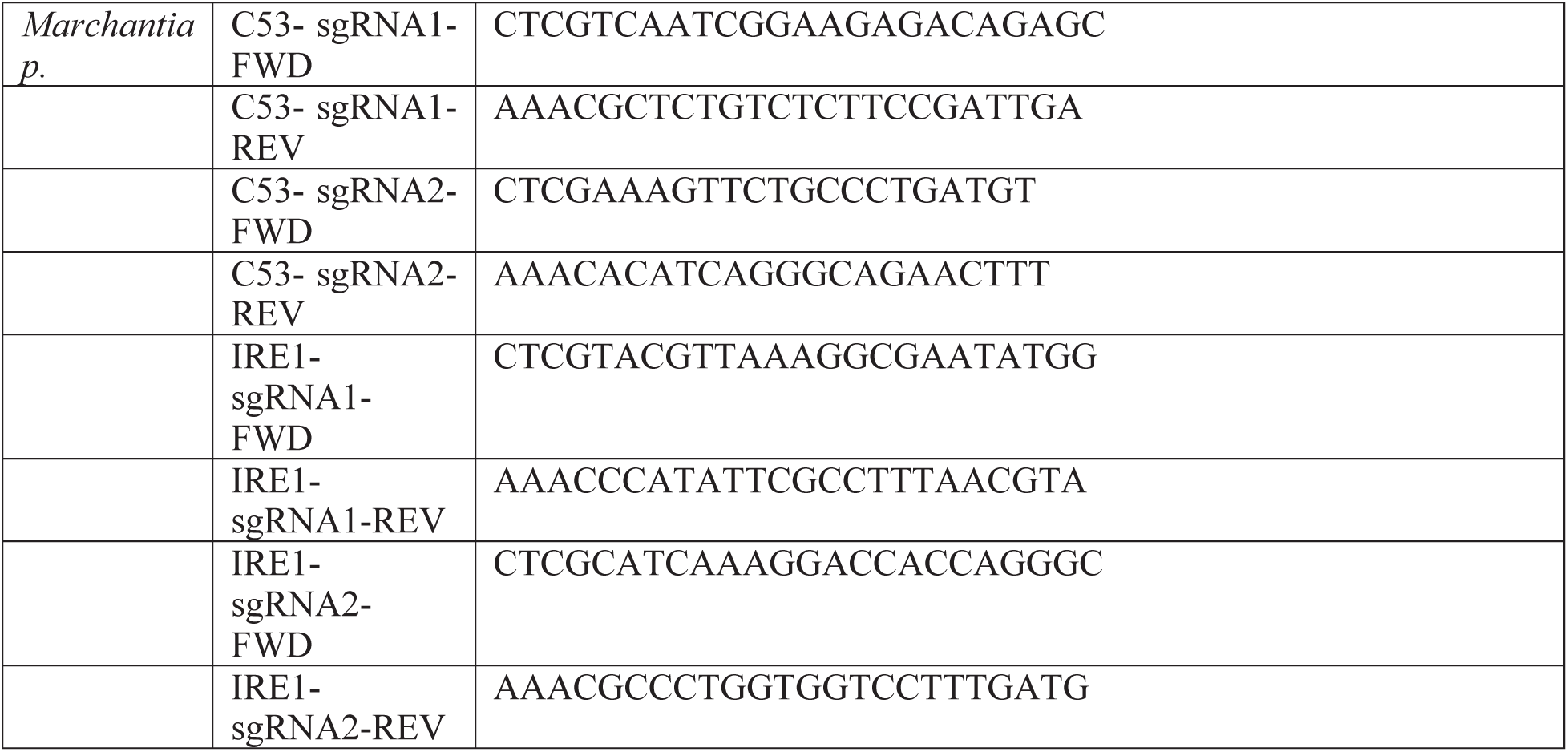

### Antibodies

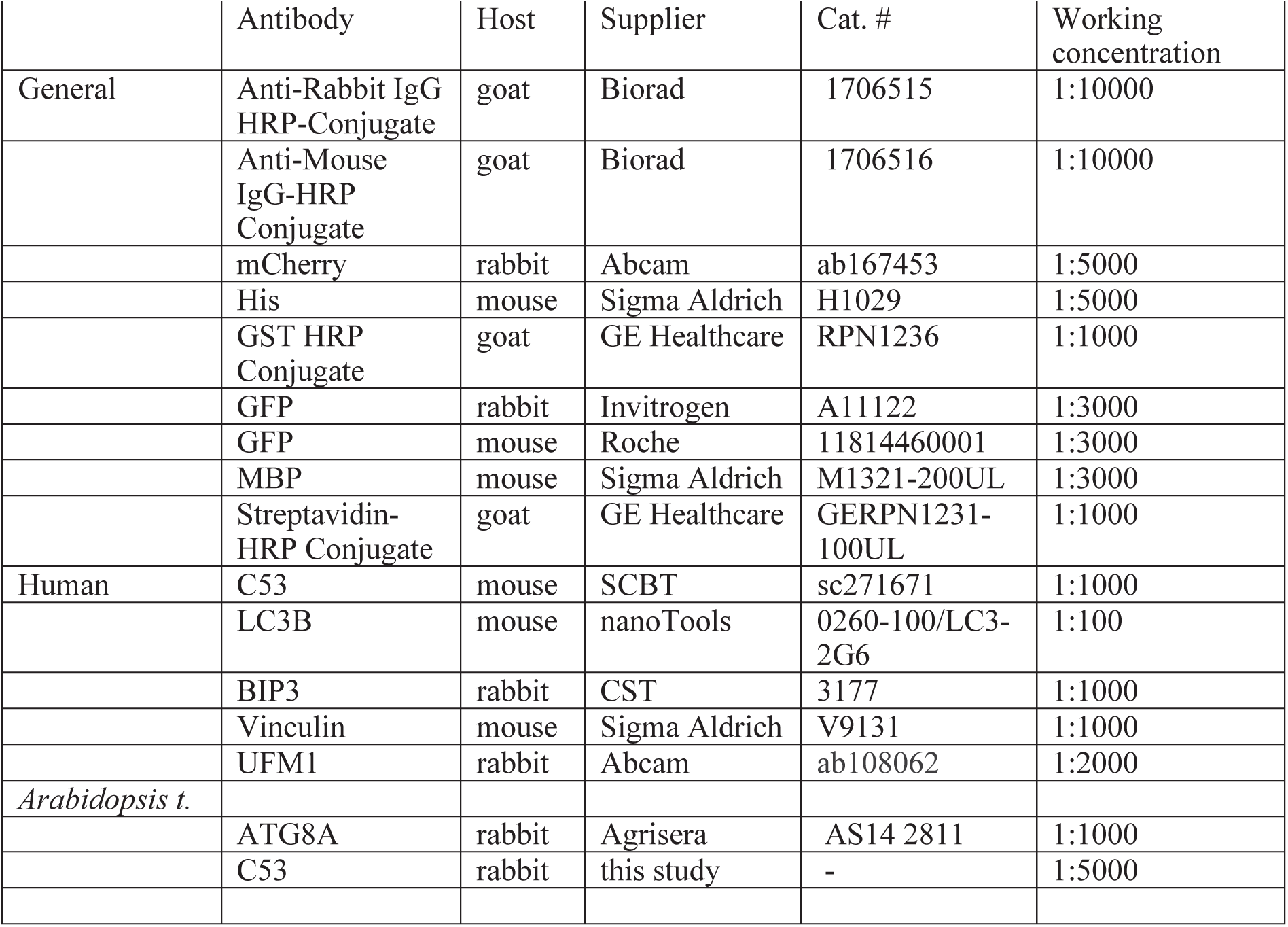

### Plasmids

| Organism | Name | Supplier | Cat. # | Reference |
| --- | --- | --- | --- | --- |
| *E. coli* |  |  |  |  |
|  | E. coli destination (expression) vector | This study |  |  |
|  | MBP-AtC53 | This study |  |  |
|  | GST-ATG8A | This study |  |  |
|  | GST-ATG8A<sup>LDS(YL50AA)</sup> | This study |  |  |
|  | GST-ATG8A<sup>UDS(IFV77AAA)</sup> | This study |  |  |
|  | GST-ATG8B | This study |  |  |
|  | GST-ATG8C | This study |  |  |
|  | GST-ATG8D | This study |  |  |
|  | GST-ATG8E | This study |  |  |
|  | GST-ATG8F | This study |  |  |
|  | GST-ATG8G | This study |  |  |
|  | GST-ATG8H | This study |  |  |
|  | GST-ATG8I | This study |  |  |
|  | GST-GABARAP | This study |  |  |
|  | GST-GABARAPL1 | This study |  |  |
|  | GST-GABARAPL2 | This study |  |  |
|  | GST-LC3A | This study |  |  |
|  | GST-LC3B | This study |  |  |
|  | GST-LC3C | This study |  |  |
|  | GST-GABARAP<sup>LDS(YL49AA)</sup> | This study |  |  |
|  | GST-GABARAP<sup>(P52A)</sup> | This study |  |  |
|  | GST-GABARAP<sup>(R67A)</sup> | This study |  |  |
|  | GST-GABARAP<sup>(P52A, R67A)</sup> | This study |  |  |
|  | GST-GABARAP<sup>(KK64AA)</sup> | This study |  |  |
|  | GST-MpATG8A | This study |  |  |
|  | GST-MpATG8A<sup>LDS(YL50AA)</sup> | This study |  |  |
|  | GST-MpATG8B | This study |  |  |
|  | GST-MpATG8B<sup>LDS(YL50AA)</sup> | This study |  |  |
|  | MBP-MpC53 | This study |  |  |
|  | MBP-AtC53 | This study |  |  |
|  | MBP-AtC53<sup>N-IDR(1-372)</sup> | This study |  |  |
|  | MBP-AtC53<sup>C-IDR(239-549)</sup> | This study |  |  |
|  | MBP-AtC53<sup>IDR(239-372)</sup> | This study |  |  |
|  | MBP-AtC53<sup>N-C(1-239,(KSGSGTSGSG)2,373-549)</sup> | This study |  |  |
|  | MBP-HsC53 | This study |  |  |
|  | MBP-HsC53<sup>N-IDR(1-316)</sup> | This study |  |  |
|  | MBP-HsC53<sup>C-IDR(263-506)</sup> | This study |  |  |
|  | MBP-HsC53<sup>IDR(263-316)</sup> | This study |  |  |
|  | MBP-HsC53<sup>N-C(1-262, (KSGSGTSGSG),317-506)</sup> | This study |  |  |
|  | MBP-AtC53<sup>Y304A</sup> | This study |  |  |
|  | MBP-AtC53<sup>Y304A, 1A (W276A)</sup> | This study |  |  |
|  | MBP-AtC53<sup>Y304A, 2A (W287A)</sup> | This study |  |  |
|  | MBP-AtC53<sup>Y304A, 3A (W335A)</sup> | This study |  |  |
|  | MBP-AtC53<sup>Y304A, 12A (W276A, W287A)</sup> | This study |  |  |
|  | MBP-AtC53<sup>Y304A, 13A (W276A, W335A)</sup> | This study |  |  |
|  | MBP-AtC53<sup>Y304A, 23A (W287A, W335A)</sup> | This study |  |  |
|  | MBP-AtC53<sup>Y304A, 123A (W276A, W287A, W335A)</sup> | This study |  |  |
|  | MBP-HsC53<sup>1A(W269A)</sup> | This study |  |  |
|  | MBP-HsC53<sup>2A(W294A)</sup> | This study |  |  |
|  | MBP-HsC53<sup>3A(W312A)</sup> | This study |  |  |
|  | MBP-HsC53<sup>12A(W269A, W294A)</sup> | This study |  |  |
|  | MBP-HsC53<sup>13A(W269A, W312A)</sup> | This study |  |  |
|  | MBP-HsC53<sup>23A(W294A, W312A)</sup> | This study |  |  |
|  | MBP-HsC53<sup>123A(W269A, W294A, W312A)</sup> | This study |  |  |
|  | MBP-AtC53<sup>IDR sAIM (Y304A, W276A, W287A, W335A)</sup> | This study |  |  |
|  | MBP-HsC53<sup>IDR sAIM(W269A, W294A, W312A)</sup> | This study |  |  |
|  | GST-AtUFL1 | This study |  |  |
|  | GST-AtC53 | This study |  |  |
|  | MBP | This study |  |  |
|  | Ts-ATG8A | This study |  |  |
|  | Ts-AtDDRKG1<sup>(24-298)</sup> | This study |  |  |
|  | Ts-AtC53 | This study |  |  |
|  | HIS6-ATG8A | This study |  |  |
|  | HIS6-GABARAP | This study |  |  |
|  | HIS6-AtC53 | This study |  |  |
|  | HIS6-AtC53<sup>sAIM (Y304A, W276A, W287A, W335A)</sup> | This study |  |  |
|  | HIS6-HsC53 | This study |  |  |
|  | HIS6-HsC53<sup>sAIM(W269A, W294A, W312A)</sup> | This study |  |  |
|  | GST | This study |  |  |
| Plant |  |  |  |  |
|  | pUbi::C53-mCherry | this study |  |  |
|  | pUbi::C53-GFP | this study |  |  |
|  | pUbi::C53<sup>sAIM(W276A, W287A, Y304A, W335A)</sup>-GFP | this study |  |  |
|  | pUbi::UFL1-GFP | this study |  |  |
|  | pUbi::DDRKG1-GFP | this study |  |  |
|  | pUbi::C53-GFP | this study |  |  |
|  | pUbi::mCherry-ATG8A | this study |  |  |
|  | pUbi::GFP-ATG8A | this study |  |  |
|  | pUbi::GFP-ATG8B | this study |  |  |
|  | pUbi::GFP-ATG8C | this study |  |  |
|  | pUbi::GFP-ATG8D | this study |  |  |
|  | pUbi::GFP-ATG8E | this study |  |  |
|  | pUbi::GFP-ATG8F | this study |  |  |
|  | pUbi::GFP-ATG8G | this study |  |  |
|  | pUbi::GFP-ATG8H | this study |  |  |
|  | pUbi::GFP-ATG8I | this study |  |  |
| Human Cell Culture |  |  |  |  |
|  | psPAX2 | Addgene | 12260 | Didier Trono |
|  | pMD2.G | Addgene | 12259 | Didier Trono |
|  | C53 shRNA in pLKO1 | Honglin Li | Gift | Wu *et al. Cell Res* (2013). |
|  | peGFP(N2)-HsC53-GFP | This study |  |  |
|  | peGFP(N2)-AtC53-GFP | This study |  |  |
|  | peGFP(N2)-HsC53<sup>sAIM</sup>-GFP | This study |  |  |
|  | peGFP(N2)-AtC53 | This study |  |  |
|  | pmCherry(N2)-HsC53-mCherry | This study |  |  |
|  | pmCherry-GABARAP-mCherry | Fumiyo Ikeda | Gift |  |
|  | mRFP-LAMTOR1 | Sascha Martens | Gift |  |
|  | ER-K20 | Addgene | 133861 | Wang *et al.* Cell Res. (2020) |
|  | ERAD-C (pEGFP-GFP:CFTR<sub>ΔF508</sub>) | Ron R. Kopito | Gift | Leto *et al.* Mol. Cell (2019) |
|  | ERAD-L (pcDNA3-NHK-GFP) | Ron R. Kopito | Gift | Leto *et al.* Mol. Cell (2019) |
|  | ERAD-M (pMCB497-pTRE-INSIG1-GFP) | Ron R. Kopito | Gift | Leto *et al.* Mol. Cell (2019) |
|  | pcDNA3-Erdj3-GFP-3Gly | Maya Schuldiner | Gift | Ast *et al.* Cell (2016) |

### General Chemicals

| Name | Supplier | Cat. # |
| --- | --- | --- |
| Vectashield | Vector | H-1000-10 |
| GeneJuice | Sigma Aldrich Aldrich | 70967 |
| Phenolred | Sigma Aldrich Aldrich | P3252 |
| Polybrene | Gift from Fumiyo Ikeda |  |
| 5x Transfer Buffer | BioRad |  |
| Acrylamide |  |  |
| Glycerol (87%) | Applichem | A0970 |
| Glass beads (0.75-1mm) | Roth | A554.1 |
| Glass beads (1.7-2.1mm) | Roth | A556.1 |
| Glass beads (2.85-3.45mm) | Roth | A557.1 |
| IGEPAL CA-630 | Sigma Aldrich | I8896-50ml |
| Magnesium Chloride | Sigma Aldrich | M8266-100G |
| Precellys steel beads 2.8mm | peqlab |  |
| Tris(2-carboxyethyl)-phosphine hydrochloride solution 0.5 M pH7 | Sigma Aldrich | 646547-10X1ML |
| Acrylamide/Bis-acrylamide, 30% solution, 37.5:1 | Sigma Aldrich |  |
| Bovine Serum Albumine | Sigma Aldrich |  |
| Glycerol (87%) | Applichem | A0970 |
| EDTA-free cOmplete Protease inhibitor Cocktail | Roche | 4693159001 |
| Tween-20 | Sigma Aldrich |  |
| TBS | In-house facility |  |
| DMSO | Sigma Aldrich |  |
| Isopropanol | Sigma Aldrich |  |
| 10x Transfer Buffer (wet) | In-house facility |  |
| pH Stripes |  |  |
| NaN<sub>3</sub> |  |  |
| NaOH |  |  |
| Ethanol |  |  |
| Naphthol Blue (Amidoblack) | Sigma Aldrich | N3393-25G |
| Acetic Acid |  |  |
| Methanol |  |  |
| ZnCl<sub>2</sub> | Gift from Fumiyo Ikeda |  |
| Triton X-100 |  |  |
| NaF |  |  |
| EGTA |  |  |
| HEPES | In-house facility |  |
| EDTA | Sigma Aldrich | E6760-100G |
| NaCl (sol.) | In-house facility |  |
| Tris Buffers | In-house facility |  |
| Bromophenolblue |  |  |
| DTT | Sigma Aldrich | 43815 |
| SDS |  |  |
| Glycerol |  |  |
| APS |  |  |
| TEMED |  |  |
| Paraformaldehyd |  |  |
| Thimerosal |  |  |
| Sodium dihydrogen phosphate |  |  |
| Sodium hydrogen phosphate |  |  |

### Inhibitors and drugs

| Name | Supplier | Cat. # |
| --- | --- | --- |
| Tunicamycin | SCBT | sc-3506 |
| Torin | SCBT | sc-396760 |
| Bafilomycin A1 | Abcam | ab120497 |
| 4μ8C | Sigma Aldrich | SML0949 |
| KIRA6 | MedChemExpress | HY-19708 |
| Anisomycin (ANS) | Sigma Aldrich | A5862-0.5ml |
| DTT | Sigma Aldrich | 43815 |
| Concanamycin-A (conA) | Santa Cruz | sc-202111A |
| Cyclopiazonic acid (CPA) | Santa Cruz | sc-201510 |
| Kifunensine (kif) | Santa Cruz | sc-201364A |
| Thapsigargin (Tg) | Santa Cruz | sc-24017 |

### Media and supplements

| Name | Supplier | Cat. # |
| --- | --- | --- |
| FBS | Sigma Aldrich | F7524 |
| PBS/PBS Tissue Culture | In-house facility |  |
| LB | In-house facility |  |
| 2xTY | In-house facility |  |
| TB | In-house facility |  |
| DMEM | In-house facility |  |
| SOC | In-house facility |  |
| gamborg B5 vitamin mixture 1000X | Duchefa | G0415.0250 |
| gamborg B5 medium (microsalt mixture) | Duchefa | M0302.0025 |
| gamborg B5 medium (including vitamins) | Duchefa | G0210.0010 |
| gamborg B5 medium (basal salt mixture) | Duchefa | G0209.0050 |
| Murashige & Skoog vitamin mixture 1000X | Duchefa | M0409.0250 |
| Murashige & Skoog micro salt mixture | Duchefa | M0301.0050 |
| Murashige & Skoog macro salt mixture | Duchefa | M0305.0050 |
| Murshige & Skoog Basal salt mixture with MES | Duchefa | M0254.0050 |
| Murashige & Skoog without nitrogen | Caisson labs |  |
| MES monohydrate | Applichem | A1074 |
| Puromycin | Sigma Aldrich | P8833 |
| Ampicillin | In-house facility |  |
| Kanamycin | In-house facility |  |
| Penicillin-Streptomycin | Sigma Aldrich | P4333 |
| Trypsin | Thermo Fisher | 25300054 |
| L-Glutamine | Sigma Aldrich | G7513 |

### Affinity matrices for purification and immuno-precipitation

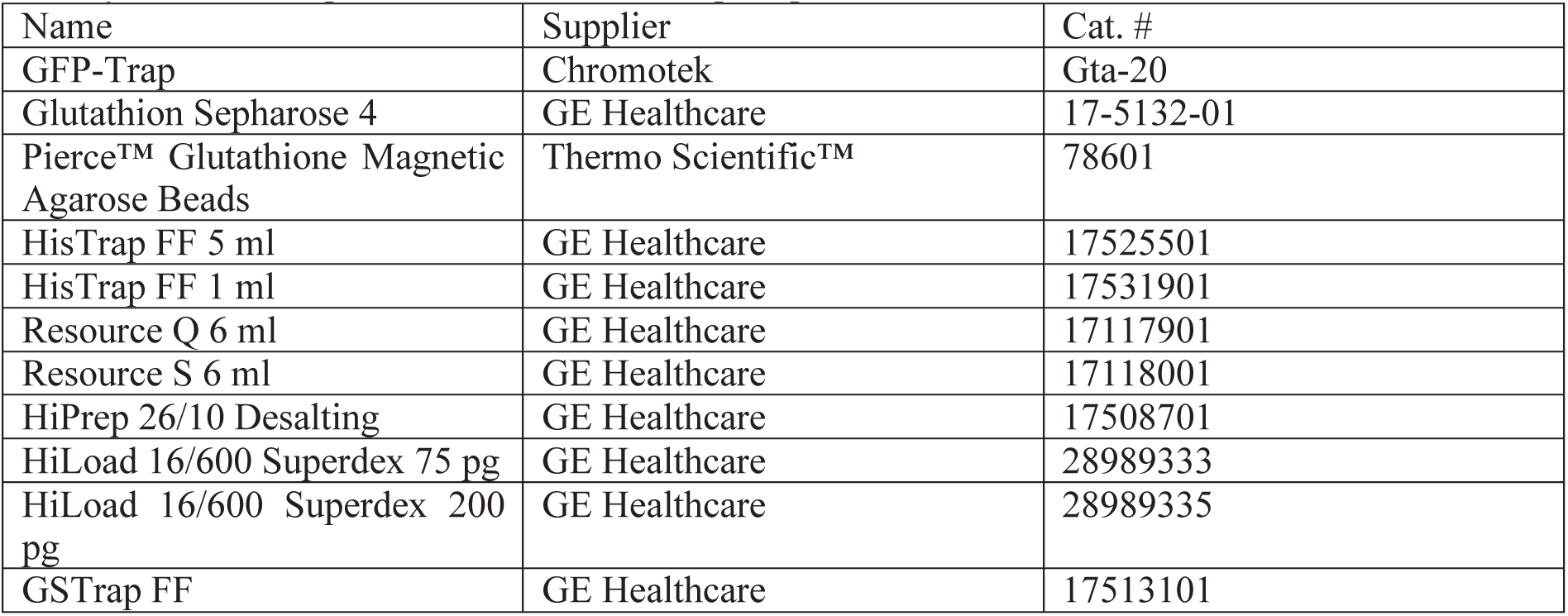

| Name | Supplier | Cat. # |
| --- | --- | --- |
| GFP-Trap | Chromotek | Gta-20 |
| Glutathion Sepharose 4 | GE Healthcare | 17-5132-01 |
| Pierce™ Glutathione Magnetic Agarose Beads | Thermo Scientific™ | 78601 |
| HisTrap FF 5 ml | GE Healthcare | 17525501 |
| HisTrap FF 1 ml | GE Healthcare | 17531901 |
| Resource Q 6 ml | GE Healthcare | 17117901 |
| Resource S 6 ml | GE Healthcare | 17118001 |
| HiPrep 26/10 Desalting | GE Healthcare | 17508701 |
| HiLoad 16/600 Superdex 75 pg | GE Healthcare | 28989333 |
| HiLoad 16/600 Superdex 200 pg | GE Healthcare | 28989335 |
| GSTrap FF | GE Healthcare | 17513101 |

### Software

| Name | Company | Application |
| --- | --- | --- |
| CLC main work bench 7 | Qiagen | Cloning |
| Zen Software | Carl Zeiss | Microscopy |
| Image J (Fiji) | NIH | Image Quantification |
| Prism 8 | Graph Pad | Statistics |
| Image Lab | BioRad | Western Blot Analysis |
| Adobe Illustrator 2020 | Adobe Inc. | Graphics editing |
| RStudio 1.2.5019 | RStudio, Inc | Graph plotting |

